# Identification of the Branching Order within the Kingdom *Bamfordvirae*

**DOI:** 10.1101/2022.05.25.493380

**Authors:** Armen Kotsinyan, Harutyun Sahakyan, Hovakim Zakaryan

## Abstract

The kingdom *Bamfordvirae* comprises the majority of the realm *Varidnaviria* and, according to the 2021 release of Virus Taxonomy by the International Committee on Taxonomy of Viruses, consists of the phyla *Nucleocytoviricota* and *Preplasmiviricota*. There are several fundamental unresolved issues related to the evolution of *Bamfordvirae*. These are questions concerning *Bamfordvirae* taxonomy including the branching order of *Nucleocytoviricota* and the question of the monophyly of *Preplasmiviricota*. Here, based on the analyses of the individual core protein phylogenies, supertree, concatenated trees, dendrograms, as well as superdendrogram, we have refined the branching order of major groups within phylum *Nucleocytoviricota* using the rooting of the entire phylum on the cellular outgroups. These efforts resulted in several major changes in *Bamfordvirae* phylogeny. In particular, we showed that *Nucleocytoviricota* consists of two sister clades, consisting of *Phycodnaviridae* sensu lato on the one hand and *Mimiviridae* sensu lato, *Iridoviridae*/*Ascoviridae*, *Marseilleviridae*, pithoviruses including *Cedratvirus*, *Solumvirus*, *Solivirus*, and *Orpheovirus*, Mininucleoviridae, *Asfarviridae* sensu lato, and *Poxviridae* on the other hand. According to our data, *Asfarviridae* sensu lato and *Poxviridae* have likely originated from within the class *Megaviricetes*. We gave evidence for polyphyly of the phylum *Preplasmiviricota* and argued for a transfer of the families *Lavidaviridae*, *Adintoviridae,* and *Adenoviridae* from the phylum *Preplasmiviricota* into the phylum *Nucleocytoviricota*. We also argued for the origin of the *Nucleocytoviricota* from small prokaryotic viruses and gave arguments against the origin of *Nucleocytoviricota* from the *Adintoviridae*/Polinton-like viruses.

**Importance:** The monophyly of *Varidnaviria*, consisting of the *Bamfordvirae* and *Helvetiavirae* kingdoms, remains a matter of debate. To confirm or refute the monophyly of *Varidnaviria*, it is important to identify the sources of origin and phylogenetic relationships common to all *Varidnaviria* genes, such as the FtsK-HerA superfamily ATPases, major and minor capsid proteins. However, such studies are relevant only if the direction of evolution of both kingdoms was firmly established. Thus, our work, by clarifying the phylogenetic relationships within the *Bamfordvirae* kingdom, provided a solid basis for studying the evolutionary relationships of this kingdom with a vast diversity of supposedly related viruses outside of *Bamfordvirae*, including *Helvetiavirae* (*Varidnaviria*).

## Introduction

The phylum *Nucleocytoviricota*, also known as nucleocytoplasmic large DNA viruses (NCLDV) comprises a group of large dsDNA viruses with presumably monophyletic origin (1, 2). Despite the official acceptance of this phylum in the ICTV Taxonomy, there are several unresolved problems associated with the origin and evolution of *Nucleocytoviricota*.

Two competing hypotheses have been proposed for the origin of NCLDV. According to the first NCLDV originated from a specific group of bacteriophages (2, 3). Alternatively, NCLDV originated from Polintons-like viruses, which, in turn, evolved from a tectivirus-like ancestor (4, 5). In both cases, it is believed that the NCLDV ancestral genome encoded only a few dozen genes, and then the ancestral genome rapidly expanded and diversified via gene duplications and horizontal gene transfer (HGT). Genomes of *Bamfordvirae* encode the FtsK-HerA superfamily ATPases, double jelly roll major capsid proteins (DJR MCP), as well as minor capsid proteins (5, 6). Moreover, the genomes of *Nucleocytoviricota* and the genomes of many *Preplasmiviricota*, in addition to these genes, also encode the Family B DNA polymerase (PolB) and D5-like helicase-primase genes (5). Excluding the major and minor capsid protein genes, the rest of the mentioned genes have apparent cellular homologs at the level of amino acid sequence homology (2). Therefore, the construction of phylogenetic trees of proteins shared by *Bamfordvirae* using cellular homologs could tip the scales in favor of one of the hypotheses.

NCLDV shares 20 conserved (core) orthologous genes (7). Some core genes also have cellular homologs. Unfortunately, they have been used only in a handful of studies. When cellular homologs were included in the analysis, trees for different genes were showing conflicting topologies of branching order of NCLDV (8). The “rooted” trees presented in the literature tend to be rooted conditionally, based on assumptions about the branching order of clades under study, but not with help of cellular outgroups from which core genes were transferred (9–11). As far as we know there are no studies, investigating the influence of the taxonomic composition of outgroups on the topology of NCLDV core proteins. In other words, the branching order of *Nucleocytoviricota* is not determined and remains a subject of discussion.

Genomes of several recently discovered viruses, such as *Pleurochrysis sp. endemic virus unk*, *Bovine rumen MELD virus*, *Sheep rumen MELD virus*, *Pleurochrysis sp. endemic virus 1a*, *Pleurochrysis sp. endemic virus 1b*, *Pleurochrysis sp. endemic virus 2* and highly divergent *Yaravirus brasiliensis* encode the PolB, DJR MCP and FtsK-HerA superfamily ATPase genes or at least some of these three genes, and this should be considered a good basis for classifying these viruses among *Bamfordvirae* taxa (12, 13).

Another group of replicators related to *Bamfordvirae* is mobile genetic elements (MGE) consisting of Polintons (also called Mavericks), cytoplasmic linear plasmids (CLP), and mitochondrial linear plasmids (MLP) (14–17). The common gene composition, the morphology of DJR MCPs, and phylogenies of several proteins of NCLDV, *Lavidaviridae*, Polintons, CLPs, and MLPs have been examined to identify the phylogenetic relationships between these viruses and MGEs (4, 5, 18, 19). The phylogenies of *Bamfordvirae* and MGEs, however, have been studied in isolation from the problem of the branching order of NCLDV, without full use of cellular orthologues, as well as without investigation of the impact of homoplasies on the topologies of the resulting trees (5, 18).

Phylogenetic relationships between individual homologous genes of NСLDV and their homologs from dsDNA viruses outside *Varidnaviria* also remain unclear. For instance, the possible relationships have been shown for the Family B DNA polymerases of NCLDV and *Herpesviridae* (20–22). However, the cited works contain several unresolved fragments of phylogenetic relationships among polymerases of *Eukaryota*, NCLDV, and several dsDNA viruses. According to the International Committee on Taxonomy of Viruses (ICTV) release from 2021, poxviruses and asfarviruses are combined into the class *Pokkesviricetes*, and the rest of *Nucleocytoviricota* form the class *Megaviricetes* (https://ictv.global/taxonomy/). Giving the equal taxonomic rank of *Pokkesviricetes* and *Megaviricetes* implies that the first bifurcation event of the last common ancestor (LCA) of *Nucleocytoviricota* led to the emergence of two lineages leading to the LCA of *Pokkesviricetes* and the LCA of *Megaviricetes*. The logic of dividing *Nucleocytoviricota* into the classes *Megaviricetes* and *Pokkesviricetes* flows from the greater sequence similarity within *Megaviricetes* in comparison with *Pokkesviricetes*, as well as on the topologies of previously constructed phylogenetic trees (8, 10). However, the long lengths of the branches of *Pokkesviricetes* may be associated with the accelerated evolution of this branch. This situation may lead to the masking of potential synapomorphies common to AP and some other branches, as well as to homoplasies. Furthermore, in our opinion, the topologies of the published trees do not imply the division of *Nucleocytoviricota* into the classes *Megaviricetes* and *Pokkesviricetes* (8). Similarly, the phylogenetic relationship between the phyla *Nucleocytoviricota* and *Preplasmiviricota* remains controversial (11). Thus, the position of the root of *Nucleocytoviricota* is logically justified, but it is not confidently determined and requires additional research.

Here we investigated all the above-mentioned problems related to the evolution of *Bamfordvirae* and proposed their solutions. We proposed a branching order of major groups within phylum *Nucleocytoviricota* based on the analysis of 18 individual core proteins phylogenies, supertree, 7 concatenated trees, and 15 dendrograms, as well as a superdendrogram. The vast majority of these topologies were obtained using cellular, as well as viral (the NCLDV D6/D11 helicases) homologs as outgroups. Based on the proposed branching order of *Nucleocytoviricota*, we argue for inclusion in the class *Megaviricetes* of the families *Asfarviridae* s.l. and *Poxviridae*. We give evidence for polyphyly of the phylum *Preplasmiviricota* and argue for a transfer of the families *Lavidaviridae*, *Adintoviridae*, and *Adenoviridae* into the phylum *Nucleocytoviricota*.

## Results

### The branching order of NCLDV. Phylogenies of 11 proteins (with outgroups) supported the primary split between phycodnaviruses s.l. and the rest of NCLDVs

#### The replication module. The transcription, and RNA processing proteins

Eukaryotic nuclear genomes encode four homologs of PolB: pol α (the main component of the primase complex), pol δ (leading and lagging strand synthesis), pol ε (main replicative polymerase), and pol ζ (involved in translesion synthesis) (23–27). Archaeal genomes encode at least ten homologs of PolB (B1-B10), while bacterial genomes encode at least three homologs of PolB (22). Eukaryotic nuclear genomes also encode three, and plant genomes encode five homologs of each of the two largest subunits of DNA-directed/dependent RNA polymerase (RNAP) (28–33). PolB and the largest two subunits of RNAP can be used for establishing the phylogenetic relationships among the NCLDV clades since they are universal markers. For further purposes, we needed the rooted trees. Preferably, trees that have been rooted in an outgroup. It has been shown, however, that the outgroup composition may affect the topological relationships between ingroup clades (34–37). To account for the possibility of topological violations between NCLDV and outgroups, we first examined the phylogenetic relationships of the PolB and RNAP cellular orthologues.

Hereinafter, all PolB alignments refer to a fragment including the Exonuclease, N-terminal, and DNA polymerase domains, starting from the active site motif Exo I and including the MotifC (22). The topology of the cellular PolB tree confirmed the close relationships of the eukaryotic pol δ, pol ζ, and archaeal PolB3 polymerases, as well as the sister relationship between the eukaryotic pol α clade and the joint pol δ + pol ζ + PolB3 clade (Fig. S1, Align. S1, Data S1) (22, 38–39). The second PolB clade included archaeal PolB1, PolB2 and eukaryotic pol ε sequences and corresponded to the Pol-B1 + Pol-B2 + pol ε + *Candidatus Heimdallarchaeota* archaeon LC_3 (HeimC3_00430) clade (39).

On the RNAP α subunit tree, we observed a sister relationship between the RPA1 and RPB1 clades (Fig. S2, Align. S2, Data S2). In contrast, the RNAP β subunit tree showed the sister relationships between the RPA2 and RPC2 subunits (Fig. S3, Align. S3, Data S3), which is consistent with previously published results (40).

The inclusion of the PolB, RNAP α, and β subunits of NCLDV in the analysis showed the sister relationship of the two clades within NCLDV. The first clade consisted of phycodnaviruses s.l., and the second clade consisted of all remaining NCLDVs (Fig. 1, Figs. S4-S5, Align. S4-S6, Data S4-S6).

**Fig. 1.**
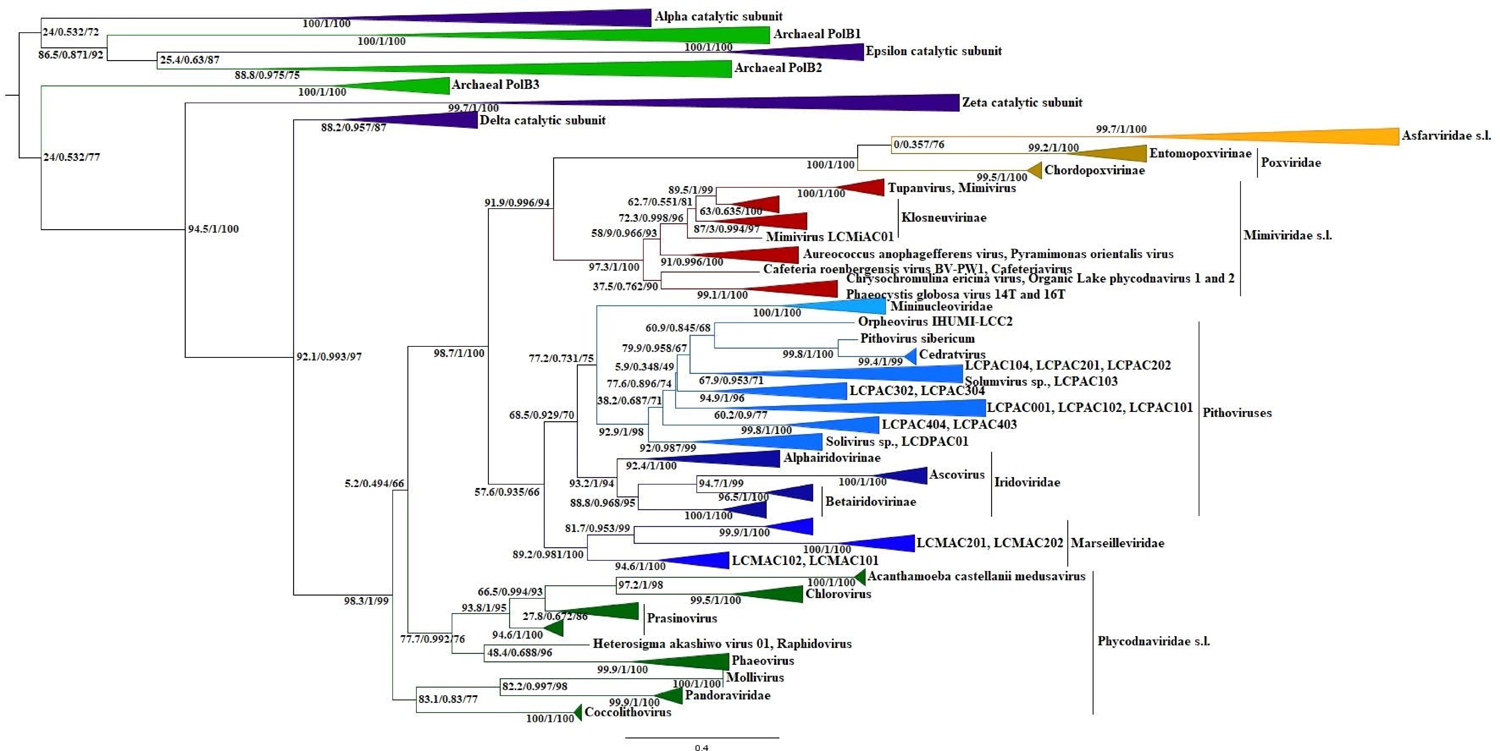
Maximum likelihood phylogenetic tree of the archaeal PolB1, PolB2, PolB3, NCLDV Family B DNA polymerases, as well as the eukaryotic pol α, pol δ, pol ε, and pol ζ catalytic subunits. The tree was constructed using the IQ TREE program (version 1.6.12) under the LG+F+R7 model and rooted according to the topology of Fig. S1, as well as in published data (22, 39). The corresponding alignment of the Exo I – MotifC fragment was built using the MUSCLE algorithm. Statistical support of the partitions, measured by SH-aLRT branch tests (1,000 replicates), approximate Bayes, and ultrafast bootstrap (1,000 replicates), is shown at the nodes. Scale bars represent the number of amino acid substitutions per site.

The origin and topological position of the *Asfarviridae* s.l. and *Poxviridae* (the AP clade) PolBs is a matter of debate and is of great systematic importance. To clarify the topological position of PolBs of asfarviruses and poxviruses, we aligned the cellular and viral polymerases and only after that added the PolB sequences of the AP clade to the alignment. We proceeded from the assumption. The AP clade PolBs will be localized outside of NCLDV if they are of non-NCLDV origin. However, polymerases of the AP clade have been found within the NCLDV clade (Fig. 1). We built the mTM-align generated multiple protein structure alignment (41, 42) consisting of the eukaryotic pol α, pol ζ, pol δ, pol ε, archaeal PolB1, PolB2, PolB3, and the AP clade predictions (Aligns. S7 (raw), S8 (trimmed)). Based on the structural alignment S8 tree showed that the AP, PolB3, pol δ, and pol ζ sequences are closer to each other than to the rest of the sequences (Data S7). Then, we selected the 851 PolB metagenomic sequences that did not have gaps in the constant vertical blocks from Aylward et al. (43), added 16 metagenomic sequences downloaded from the WGS metagenomic projects (env_nr) database, and using the alignment S8 as a template aligned metagenomic sequences with the homologs of NCLDV, PolB3, pol δ, and pol ζ using the MUSCLE v5 software (44). It is important to note, that the sequences of the NCLDV families (*Asfarviridae* s.l., *Poxviridae*, *Iridoviridae*, etc.), as well as cellular polymerases (pol δ, pol ζ, PolB3), were aligned separately before being combined and aligned. Based on this alignment, we constructed the following trees consisting of a) PolB3, pol δ, pol ζ, and viral (metagenomic and identified) homologs (Align. S9, Data S8), b) pol δ and viral homologs (Fig. 2, Align. S10, Data S9), and c) pol δ and viral homologs (without the metagenomic and identified *Mimiviridae* s.l. sequences) (Align. S11, Data S10). The topology presented in Data S8 and Fig. 2 showed that phycodnaviruses s.l. was the basal group of viruses to crown clades of NCLDV. In turn, the joint *Mimiviridae* s.l. + AP and MAPIM clades demonstrated a sister relationship. However, a) the AP clade was localized sisterly to *Mimiviridae* s.l. in Figure 2, whereas a sister relationship was detected between asfarviruses s.l. and mimiviruses s.l in Data S8, while poxviruses formed a sister clade to the joint *Mimiviridae* s.l. + *Asfarviridae* s.l. clade. b). In Figure 2 the pol δ/*Herpesviridae*-related clade was localized sisterly to the joint *Mimiviridae* s.l. + MAPIM + AP clade (hereinafter the MAPIMAPM clade), whereas in Data S8 this clade formed a sister clade to MAPIM. c). The *Iridoviridae*-related clade was localized basal to the MAPIM families in Figure 2, whereas this clade was observed within the MAPIM clade in Data S8. The removal of mimiviruses s.l. (including metagenomic sequences) from the alignment S10 did not change the topological localization of the AP clade, which speaks in favor of the absence or minimal effect of homoplasies on the position of the AP clade (Data S10). Thus, the results spoke in favor of the origin of the *Asfarviridae* s.l. and *Poxviridae* polymerases from within NCLDV, in contrast to previously published data (22). However, this assumption is more controversial for poxviruses than for asfarviruses (see below).

**Fig. 2.**
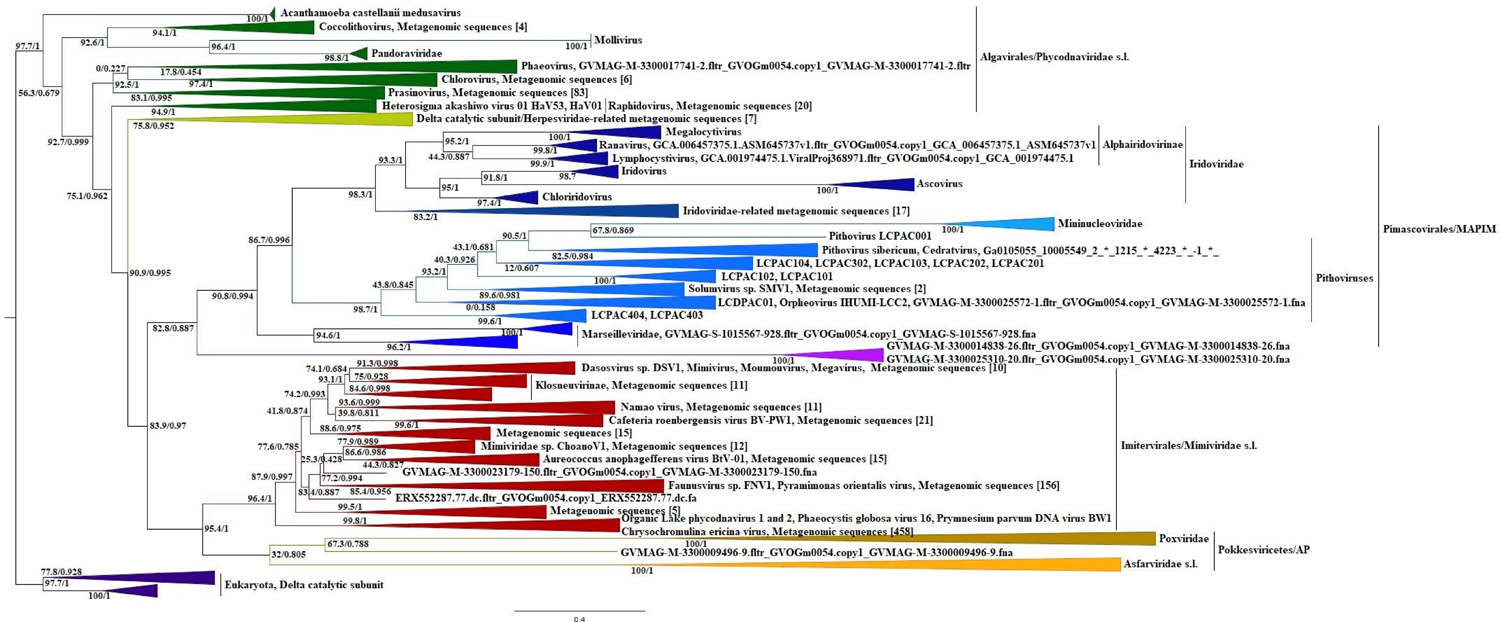
Maximum likelihood phylogenetic tree of the eukaryotic δ catalytic subunits, NCLDV, and viral metagenomic Family B DNA polymerases. The tree was constructed using the IQ TREE program (version 1.6.12) under the LG+F+I+R4 model and rooted according to the topologies of Fig. S1 and Fig. 1, as well as in published data (22, 39). The corresponding alignment of the Exo I – MotifC fragment was built using the MUSCLE v5 software under the permutation none, perturbation default parameters, and using the alignment S8 as a template. Square brackets indicate the number of metagenomic sequences belonging to a given clade. Statistical support of the partitions, measured by SH-aLRT branch tests (1,000 replicates) and approximate Bayes, is shown at the nodes. Scale bars represent the number of amino acid substitutions per site.

The branches corresponding to PolB, RNAP α, and RNAP β of NCLDV are long and potentially may contain many character states uniting the NCLDV clades with outgroup clades (homoplasies), especially with the sister clades to NCLDV. This can lead to a false topological position of NCLDV among the cellular clades, as well as to false topological relationships of viral clades within NCLDV. To minimize the possibility of the influence of homoplasies, we removed the pol δ, RPB1, and RPB2 sequences from the corresponding alignments (Align. S4-S6) and reconstructed the trees (Figs. S6-S8, Aligns. S12-S14, Data S11-S13). The reconstruction showed the invariability of topological relationships among the NCLDV clades, as well as among NCLDV and the cellular clades, which suggests that homoplasies were unlikely.

Changing the composition of outgroups is a reliable way to check the influence of the composition of outgroups on the topology of ingroups (45). We built the alignments of the pol δ, RNAP α and β subunits using the MUSCLE and MAFFT algorithms, in which we retained only the sequences of NCLDV and corresponding eukaryotic sister clades and reconstructed the trees (Figs. S9-S14, Aligns. S15-S20, Data S14-S19). Next, we completely removed all outgroups and reconstructed the trees (Figs.S15-S17, Aligns. S21-S23, Data S20-S22). The resulting topologies showed that regardless of the taxonomic composition of outgroups the relationships between clades within NCLDV remain unchanged.

Thus, regardless of the scenario whether the PolB, RNAP α, and β genes of NCLDV were captured from an ancestral lineage leading to modern eukaryotes or from now extinct protoeukaryotic lineages, the identified order of the first two bifurcation events of NCLDV remained unchanged. Therefore, the revealed branching order may reflect the true evolutionary trajectory of NCLDV. Additional confidence in the results was provided by a comparison of earlier results with ours, which confirmed the close relationship between the eukaryotic pol δ catalytic subunits and NCLDV polymerases (20–22).

Unlike other NCLDVs, phycodnaviruses s.l. encodes two different ORFs for the helicase-primase (SFIII helicase/D5-like primase) gene. For instance, in the *Paramecium bursaria Chlorella virus NY2A* (NC_009898) genome, the SFIII helicase localizes in the region 260150…262114 nucleotides, while the D5-like primase localizes in the region 265781…267112 nucleotides. This situation does not exclude the possibility of different evolutionary trajectories of primase and helicase of phycodnaviruses s.l., of which only one can coincide with the helicase or primase trajectory of the rest of NCLDVs. Consequently, the resulting phylogenetic tree constructed using the concatenated sequences of *Phycodnaviridae* s.l. may not reflect the true evolutionary trajectory of NCLDV (for instance, in Figure 1C (8), see below). To analyze two potentially different trajectories of D5-like primases and SFIII helicases of phycodnaviruses s.l., we examined them as two separate proteins. Indeed, the topological relationships of the D5-like clades of NCLDV, obtained using outgroups, as well as without, repeated the trajectories obtained for polymerases, that is, phycodnaviruses s.l. formed a sister clade to the rest of NCLDVs, and mimiviruses s.l. formed a sister clade to MAPIM (Figs. S18-S20, Aligns. S24-26, Data S23-S25). At the same time, the topology of SFIII helicases obtained using of bacterial (phage/plasmid primase, P4 family) outgroup showed affinity between the phage/plasmid primase sequences of *Gammaproteobacteria* and homologs of *Phycodnaviridae* s.l. and separated the *Phycodnaviridae* s.l. clade from the rest of the NCLDVs (Fig. S21, Align. S27, Data S26). In contrast, the phylogenetic trajectories of the D5-like primases and SFIII helicases of the *Mimiviridae* s.l. and joint MAPIM + *Asfarviridae* or *Chordopoxvirinae* clades demonstrated the invariance of sister relationships (Figs. S18-S22, Align. S28, Data S27).

We identified the sequences most closely related to the poxvirus D6/D11 helicases based on preliminary constructed trees consisting of the cellular and viral helicases belonging to the superfamily II (Align. S29-S30, Data S28-S29) and constructed the separate D6/D11 phylogenetic tree (Fig. 3, Align. S31, Data S30). The resulting tree was rooted between branches containing the homologs of the ASFV D1133L and ASFV Q706L proteins, since in this case, the branch containing the ASFV D1133L homologs allowed us to show the repeatability of the branching order of NCLDV, demonstrated above via the PolB, RNAP α and β, D5-primase, and SFIII helicase topologies. That is, sister relationships were identified between the joint MAPIM + AP (hereinafter the MAPIMAP clade) and *Mimiviridae* s.l. clades, as well as between the AP and MAPIM clades. However, in contrast to the topologies mentioned above, in the case of the D6/D11 helicases, the branching order repeatability was provided not by cellular outgroups, which can potentially be problematic, but by extremely closely related homologs of NCLDV (branch containing the ASFV Q706L homologs).

**Fig. 3.**
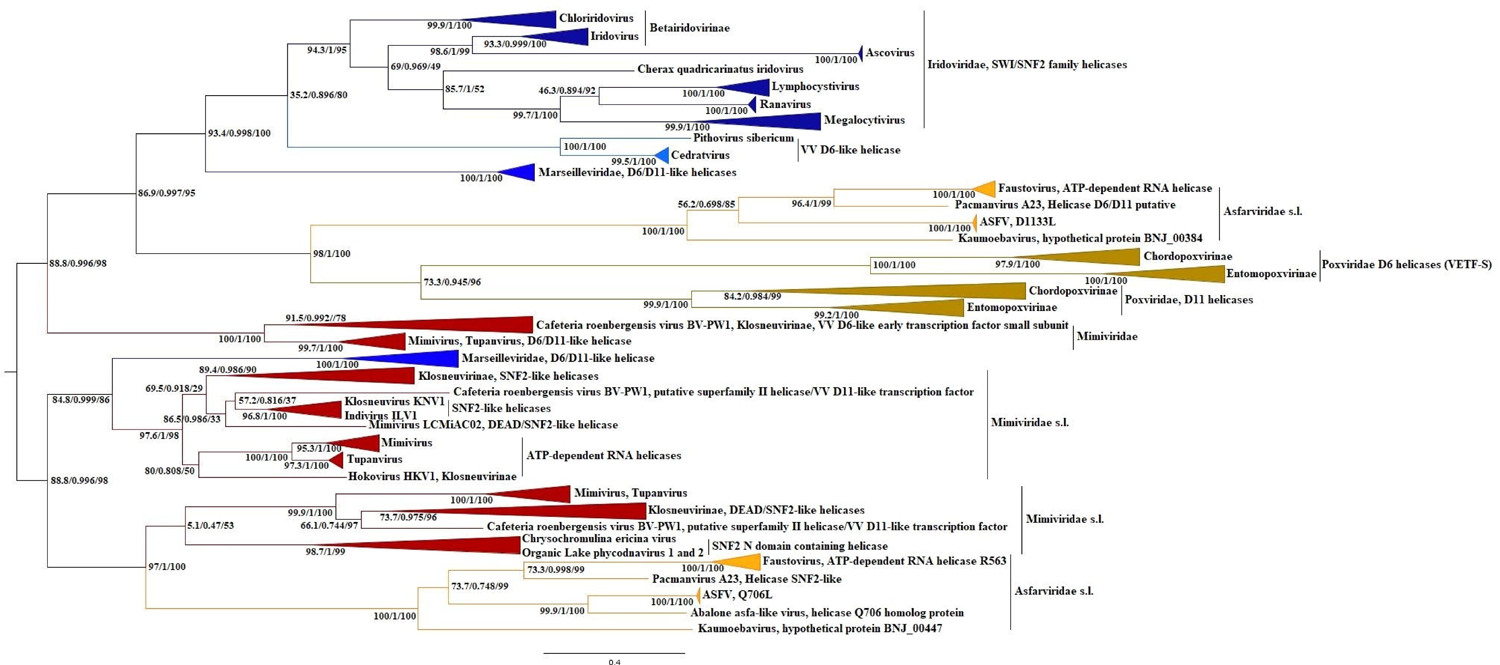
Maximum likelihood phylogenetic tree of superfamily II D6/D11-like helicases. The tree was constructed using the IQ TREE program (version 1.6.12) under the LG+F+R6 model and rooted between two clades containing the homologous sequences of *Mimiviridae* s.l., *Marseilleviridae*, and *Asfarviridae* s.l. (containing the ASFV D1133L and Q706L clades). The corresponding alignment was built using the MUSCLE algorithm. Statistical support of the partitions, measured by SH-aLRT branch tests (1,000 replicates), approximate Bayes, and ultrafast bootstrap (1,000 replicates), is shown at the nodes. Scale bars represent the number of amino acid substitutions per site.

Here we also presented the phylogenies of the remaining proteins of the replication module, as well as proteins belonging to the functional category of RNA transcription and processing. These are the AP (apurinic) endonuclease family 2 (Figs. S23-S24, Aligns S32-S33, Data S31-S32), Metallopeptidase WLM (Figs. S25-S26, Aligns. S34-35, Data S S33-S34), PCNA (Figs. S27-S28, Aligns. S36-S37, Data S35-S36), RuvC (Figs. S29-S30, Aligns. S38-S39, Data S37-38), Uracil DNA glycosylase (Figs. S31-S32, Aligns. S40-S41, Data S39-S40), Topoisomerase II (Figs. S33-S34, Aligns. S42-S43, Data S41-S42), mRNA capping enzyme (Figs, S35-S36, Aligns. S44-S45, Data S43-S44), and TFIIB (Figs. S37-S39, Aligns. S46-S48, Data S45-S47) proteins. The presented phylogenies were obtained using cellular outgroups, as well as without them. We focused on the most important bifurcations of the obtained phylogenies summarized in (Table 1). The phylogenies of all individual proteins where the gene was encoded by the genomes *Phycodnaviridae* s.l. (except for SFIII helicases (see above), as well as for Uracil DNA glycosylase, where the gene was present only in pandoraviruses and had a different fate than the rest of NCLDVs) confidently demonstrated primary bifurcation leading to the emergence of *Phycodnaviridae* s.l. and the clade consisting of the rest of NCLDVs. Phylogenies of eight proteins localized the AP clade sisterly to MAPIM, and only phylogenies of PolB (the MUSCLE/MUCLE v5 software derivatives) and Metallopeptidase WLM localized the AP clade sisterly to *Mimiviridae* s.l. Similar, phylogenies of eight proteins localized the joint AP + MAPIM clade sisterly to *Mimiviridae* s.l.

**Table 1.**
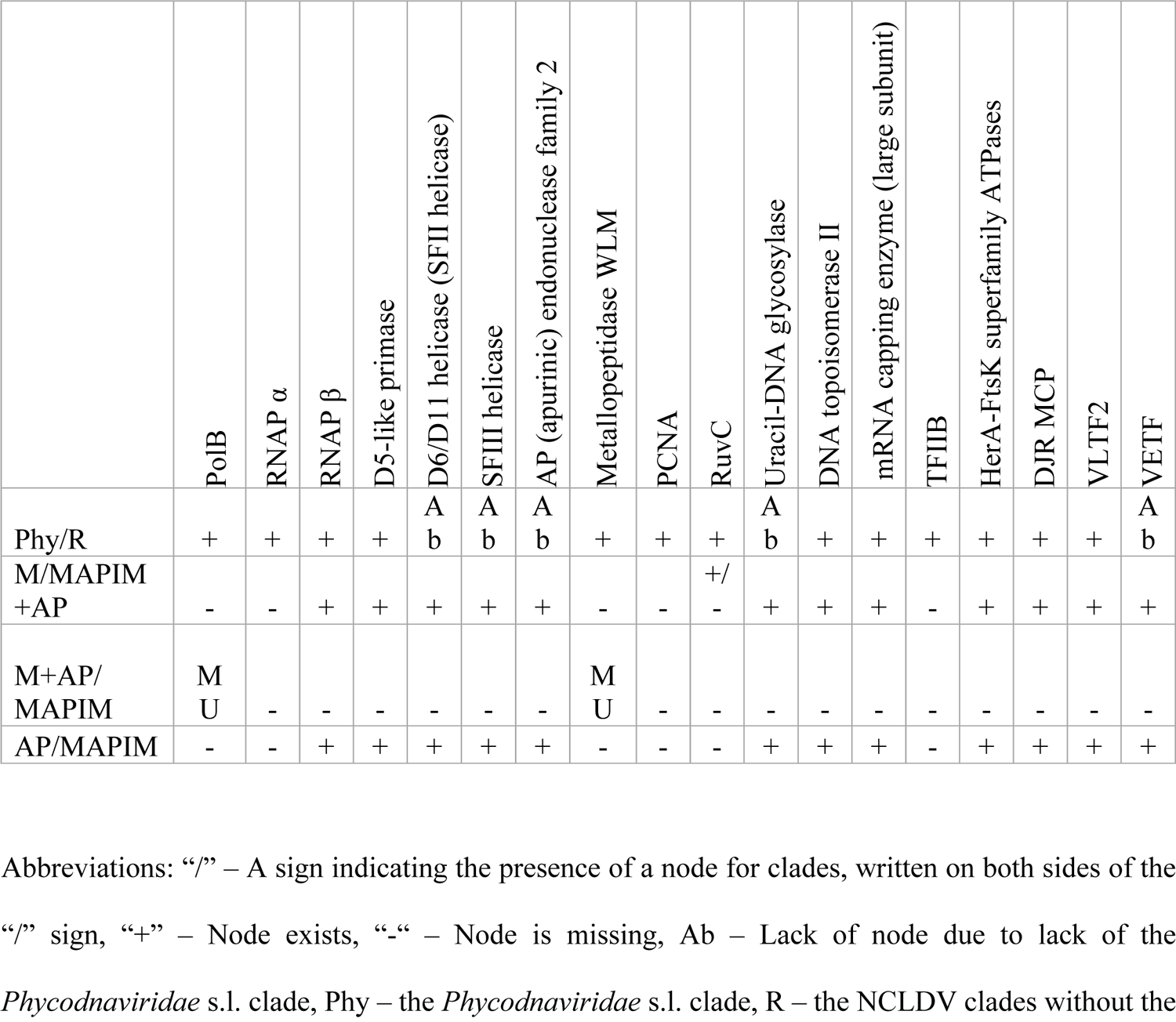
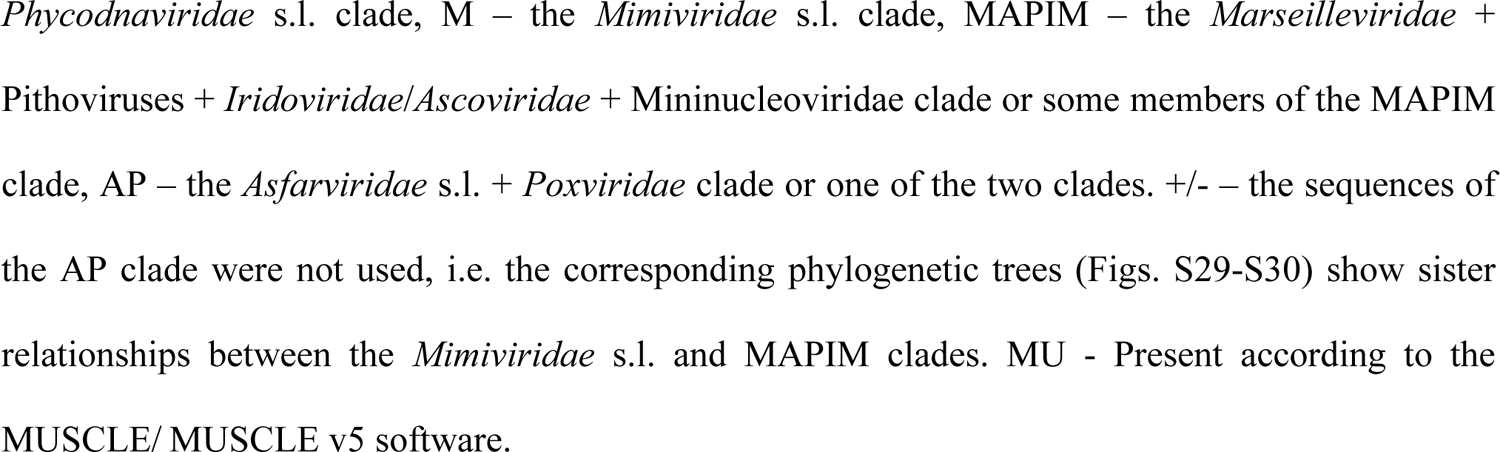
Order of bifurcations of deep nodes of the NCLDV 18 core proteins.

| | PolB | RNAP $\alpha$ | RNAP $\beta$ | D5-like primase | D6/D11 helicase (SFII helicase) | SFIII helicase | AP (apurinic) endonuclease family 2 | Metallopeptidase WLM | PCNA | RuvC | Uracil-DNA glycosylase | DNA topoisomerase II | mRNA capping enzyme (large subunit) | TFIIB | HerA-FtsK superfamily ATPases | DJR MCP | VLTF2 | VETF |
| --- | --- | --- | --- | --- | --- | --- | --- | --- | --- | --- | --- | --- | --- | --- | --- | --- | --- | --- |
| Phy/R | + | + | + | + | A<br>b | A<br>b | A<br>b | + | + | + | A<br>b | + | + | + | + | + | + | A<br>b |
| M/MAPIM<br>+AP | - | - | + | + | + | + | + | - | - | +/- | + | + | + | - | + | + | + | + |
| M+AP/<br>MAPIM | M<br>U | - | - | - | - | - | - | M<br>U | - | - | - | - | - | - | - | - | - | - |
| AP/MAPIM | - | - | + | + | + | + | + | - | - | - | + | + | + | - | + | + | + | + |

Abbreviations: “/” – A sign indicating the presence of a node for clades, written on both sides of the “/” sign, “+” – Node exists, “-” – Node is missing, Ab – Lack of node due to lack of the *Phycodnaviridae* s.l. clade, Phy – the *Phycodnaviridae* s.l. clade, R – the NCLDV clades without the

The tertiary structure of proteins is largely but not completely determined by the amino acid primary structures. The prediction of the tertiary structures is still an unsolved problem (46). Moreover, unlike the computationally expensive but attractive physical interaction program, the evolutionary program imposes structural constraints based on bioinformatics analysis of the evolutionary history of proteins (46–48). Thus, dendrograms based on a comparison of crystallographic structures and predictions are not completely phylogeny-independent evidence for the evolutionary trajectory of the clade under study. However, if the dendrogram, at least in general form, repeats the bifurcation order of phylogenetic analysis, this speaks in favor of the reliability of the obtained phylogenetic tree. Logically, dendrograms obtained via calculation of predictions of the physical interaction programs or crystallographic structures should be more reliable, since in both cases the tertiary structures and, as a result, dendrograms do not depend on bioinformatics analysis. However, the quantitative limitation of crystallographic structures and the computationally expensive physical approach limit the possibilities of comparing protein tertiary structures only to the bioinformatics approach.

We predicted the Exo I – MotifC fragment of the *Acanthamoeba castellanii medusavirus*, ASFV, *Faustovirus*, *Choristoneura biennis entomopoxvirus*, *Vaccinia virus*, *Saccharolobus solfataricus* P11 (PolB1), *Ignisphaera aggregans* (PolB2), and *Methanolobus psychrotolerans* (PolB3) polymerases, as well as the pol δ, pol ζ, pol ε, and pol α subunits of some eukaryotes using ColabFold. The resulting dendrogram based on these predictions (Data S48) showed two fundamental differences from phylogenies. Firstly, asfarviruses together with PolB3, pol ζ, pol δ, and Medusavirus formed a clade, but asfarvirus predictions localized sisterly to the joint PolB3 + pol δ + Medusavirus clade. Secondly, poxviruses were the most distant clade to the joint PolB3 + pol ζ + pol δ + Medusavirus + *Asfarviridae* clade. Meanwhile, the Z-score table (Table S1) showed that, except for themselves, asfarviruses had the highest Z-scores with Medusavirus, pol δ, and PolB3. Simultaneously, poxviruses had the highest Z-scores with asfarviruses, PolB3, Medusavirus, and pol δ. These data contradicted the topology of the dendrogram Data S48. Probably they were a consequence of the work of the DALI algorithm (49–51). Moreover, a tree consisting of the pol δ, pol ζ, PolB3, *Asfarviridae* s.l., and *Poxviridae* sequences and constructed using the MUSCLE v5 software placed the AP clade sisterly to the joint pol δ+ pol ζ clade (Align. S49, Data S49). These data supported the hypothesis of Koonin and Yutin (10), since they suggested the presence of the LCA of *Megaviricetes*, but shifted the time interval of the primary acquisition event of the protoeukaryotic PolB by the ancestral NCLDV lineage to the time before the origin of the eukaryotic pol δ and pol ζ polymerases, which in turn suggested that the ancestral PolB of *Megaviricetes* was acquired secondarily. However, this scenario conflicts with the large amount of phylogenetic data related to other proteins (see above), as well as the data presented below.

We compared the morphologies of the predicted Exo I – MotifC fragment of viral and cellular polymerases (Fig. 4A-4Z). Morphological analysis of the predicted Exo I – MotifC fragments revealed the presence of a beta-hairpin at the N-terminus of the predicted fragments in the β16-β17 region (Fig. 4A, Table S2,) among NCLDV, pol δ, pol ε, PolB2, and PolB3. However, while the beta-hairpin lengths of NCLDV and pol δ varied from 16 to 21 amino acids, the pol ε, PolB2, and PolB3 beta-hairpin lengths were 9, 10, and 12 amino acids, respectively (Fig. 4I1, Table S2, Align. S50). It was difficult to assume that this potential split, determining the NCLDV polymerases and pol δ on the one hand and the archaeal PolB2 and PolB3 and pol ε, on the other, as two sister clades, was synapomorphic, since it contradicted the phylogenies obtained by us (Fig. S1) and other authors (22). Alternatively, we should have assumed that the replacement/loss of just a few amino acids led to the appearance/disappearance or change in the hairpin length in the β16-β17 fragment. Whereas, sequences similarity and stability of beta-hairpin length among NCLDV polymerases and pol δ were more similar to synapomorphies and did not contradict the results of phylogenies. In our opinion, the discussed morphological feature spoke in favor of the common origin of the polymerases of asfarviruses and other NCLDVs, as well as δ catalytic subunits (Fig. 4B-4C, 4F-4T, 4I1). In contrast to all NCLDVs and pol δ, the predicted morphologies of the β16-β17 poxvirus fragments contained a unique feature. Within the β16-β17 fragment, the genomes of poxviruses encoded two beta hairpins (Fig. 4D-E, 4I1). The C termini of the first beta-hairpins of poxviruses coincided with the N termini of the NCLDV and pol δ beta-hairpins and overlapped with them with a length of 5-8 amino acids. However, the C termini of the first beta-hairpins of poxviruses did not either overlap or overlapped with 1-2 amino acids with the N termini of the pol ε, PolB2, and PolB3 beta-hairpins. The second beta-hairpins of poxviruses completely coincided with the middle part and C termini of the NCLDV and pol δ beta-hairpins, while with the C termini of the pol ε, PolB2, and PolB3 beta-hairpins the second beta-hairpins of poxviruses had much less similarity (Fig. 4I1). Thus, despite the insertion of 12 amino acids into the second hairpin of chordopoxviruses (Fig. 4I1), judging by the amino acid composition, the common origin of the *Poxviridae*, NCLDV, and pol δ hairpins seemed more likely than the origin of the *Poxviridae* hairpins from the PolB2, PolB3 or pol ε hairpins.

**Fig. 4.**
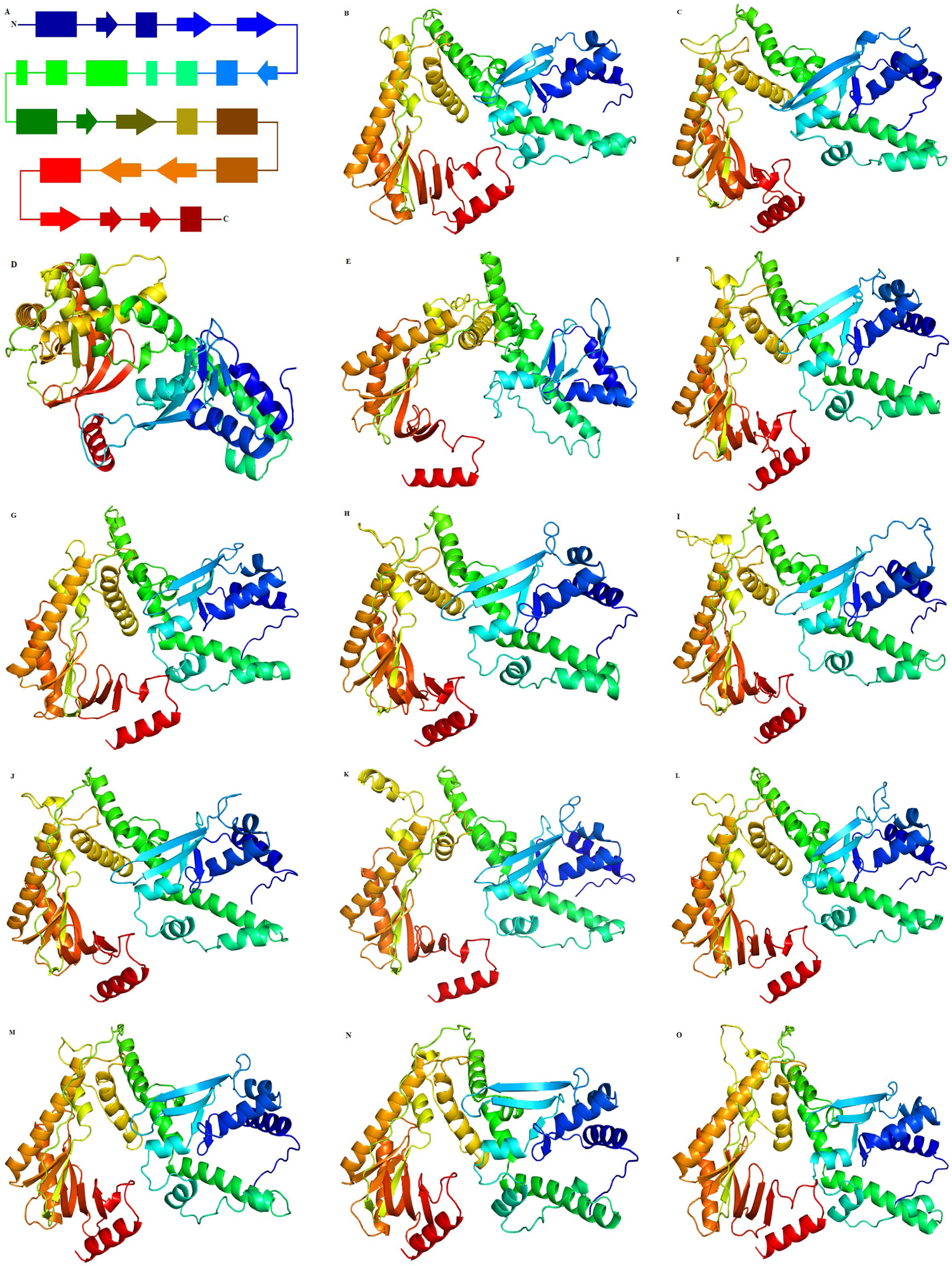
Comparison of the predicted Exo I – MotifC fragments of Family B polymerases. A. – Schematic diagram of the predicted Exo I – MotifC fragment of the δ catalytic subunit of *Saccharomyces cerevisiae* (3IAY_A). Rectangles and arrows represent α-helices and β-strands, respectively, colored using a rainbow scheme (N terminus – blue, C terminus – red). The lengths of rectangles and arrows roughly correspond to proportions of lengths of predicted α-helices and β-strands. The β16-β17 fragment, in which the beta-hairpin is located, corresponds to the second and third arrows from the N terminus (colored blue). **B** – *African swine fever virus* SPEC_57 (QGM12923), **C** – *Faustovirus* liban (QJX71044), **D** – *Yokapox virus* DakArB 4268 (YP_004821396), **E** – *Choristoneura biennis entomopoxvirus* (P30319), **F** – *Lymphocystis disease virus* - isolate China (YP_073706), **G** – *Invertebrate iridescent virus 22* IIV22 Aberystwyth (YP_009010762), **H** – *Melbournevirus* 1 (YP_009094792), **I** – *Powai lake megavirus* 1 (ANB50688), **J** – *Mimivirus LCMiAC01* (QBK88617), **K** – *Cafeteria roenbergensis virus BV-PW1* (YP_003970183), **L** – *Organic Lake phycodnavirus 1* (ADX06143), **M** – *Aureococcus anophagefferens virus* BtV-01 (YP_009052217), **N** – *Heterosigma akashiwo virus 01* HaV53 (YP_009507574), **O** – *Feldmannia irregularis virus a* FirrV-1 (YP_009665694), **P** – *Pandoravirus dulcis* Melbourne (YP_008318996), **Q** – *Paramecium bursaria Chlorella virus CVK2* K2 (BAA35142), **R** – *Acanthamoeba castellanii medusavirus* (BBI30551), **S** – *Ostreococcus tauri virus 2* (YP_004063640), **T** – *Saccharomyces cerevisiae*, DNA polymerase δ catalytic subunit (3IAY_A), **U** – *Saccharomyces cerevisiae*, DNA polymerase ζ catalytic subunit (6V93_A), **V** – *Callorhinchus milii*, DNA polymerase α catalytic subunit (AFO93960), **W** – *Saccharomyces cerevisiae*, DNA polymerase ε catalytic subunit A (6S2E_A), **X** – *Saccharolobus solfataricus* P1, PolB1 (SAI84106), **Y** – *Ignisphaera aggregans* DSM 17230, PolB2 (ADM27371), **Z** – *Methanolobus psychrotolerans*, PolB3 (WP_094229042), **A1** – *Macaca nemestrina rhadinovirus 2* Mne442N (AAF81664), **B1** – Polinton 1_XT (*Xenopus tropicalis*), **C1** – *Zea mays*, mitochondrial linear plasmid (P10582), **D1** – *Peltigera dolichorrhiza*, mitochondrial linear plasmid (YP_009316284), **E1** – *Bovine rumen MELD virus* (DAC81507), **F1** – *Coemansia reversa* NRRL 1564, cytoplasmic linear plasmid (PIA12548), **G1** – *Stylophora coral adintovirus* (DAC81741), **H1** – *Human adenovirus 54* (YP_003038603), **I1** – Alignment of the beta hairpins of NCLDV, pol δ and pol ε, as well as PolB2 and PolB3, **J1** – Alignment of the beta hairpins of ASFV SPEC_57, *Abalone asfa-like virus*, Polintons, mitochondrial and cytoplasmic linear plasmids, *Bovine rumen MELD virus*, and *Adenoviridae*. The I1 and J1 alignments were obtained by excising the β16-β17 fragment from the alignments S50 and S96, respectively. Both alignments were built using the MUSCLE v5 software. **K1** – Sheep rumen MELD virus (DAC81716), **L1** – Stylophora coral adintovirus (DAC81741).

The polymerases of asfarviruses and poxviruses were most related to the polymerases of NCLDV, pol δ, and PolB3 according to the results of phylogenies, morphology, and Z-scores. This assumption was in direct contradiction with the topology of the dendrogram S48 (Data S48). However, the topological discrepancy between the dendrogram S48 and several phylogenetic and morphological evidence suggested that phylogenies should better reflect evolutionary trajectories over the giant evolutionary distances such as the evolution of the Family B polymerases. Otherwise, we would be forced to assume extremely unlikely phylogenetic hypotheses related to the rerooting of the dendrogram. The calculation of dendrograms should be accompanied by parallel phylogenetic and morphological studies and should be limited to certain evolutionary frameworks associated with the preservation of monophyly of clades (see below).

Based on the ColabFold predictions, we calculated dendrograms of the PolB (Fig. 5 (the Exo I – MotifC fragment, 400-421 amino acids predictions), Fig. S40 (91-97 amino acids prediction), Data S50-S51), RNAP α (Fig. S41, Data S52), RNAP β (Fig. S42, Data S53), PCNA (Fig. S43, Data S54), TFIIB (Fig. S44, Data S55), Topoisomerase II (Fig. S45, Data S56), and Uracil DNA glycosylases (Fig. S46, Data S57) proteins belonging to the replication module and functional category of RNA transcription and processing. It should be noted that the PolB predictions with 91-97 amino acid length were part of the Exo I – MotifC fragment and corresponded to the MotifB (αP) - KxY (β29-αS) region (22). Z-scores and PDB25 search results for predicted structures are shown in Tables S1 and S3. The individual protein dendrograms were rooted according to the topologies of the corresponding individual protein phylogenies. The most important result, demonstrated by all dendrograms, was the repetition of the branching order of NCLDV obtained using the phylogenies of individual proteins. Phycodnaviruses either were localized basal to crown NCLDVs (PolB (Fig. 5), RNAP α (Fig. S41), PCNA (Fig. S43), Topoisomerase II (Fig. S45)) or formed a sister clade to all other NCLDVs (PolB (Fig. S40), RNAP β (Fig. S42)), TFIIB (Fig. S44)). In its turn, mimiviruses s.l. either were localized sisterly to the MAPIMAP clade (Figs. S41-S43, S46), or were localized sisterly to MAPIM (Fig. 5, Fig. S44), or were localized basal to the crown MAPIM and AP families. However, in contrast to individual phylogenies, the position of the AP clade was more variable. In Figure 5 the AP clade was localized sisterly to the joint *Mimiviridae* + MAPIM clade, whereas in Figure S40 the AP clade was localized sisterly to the MAPIM clade, which is consistent with results for the vast majority of individual phylogenies. In its turn, the topologies of the remaining dendrograms placed the AP clade within the MAPIM clade.

**Fig. 5.**
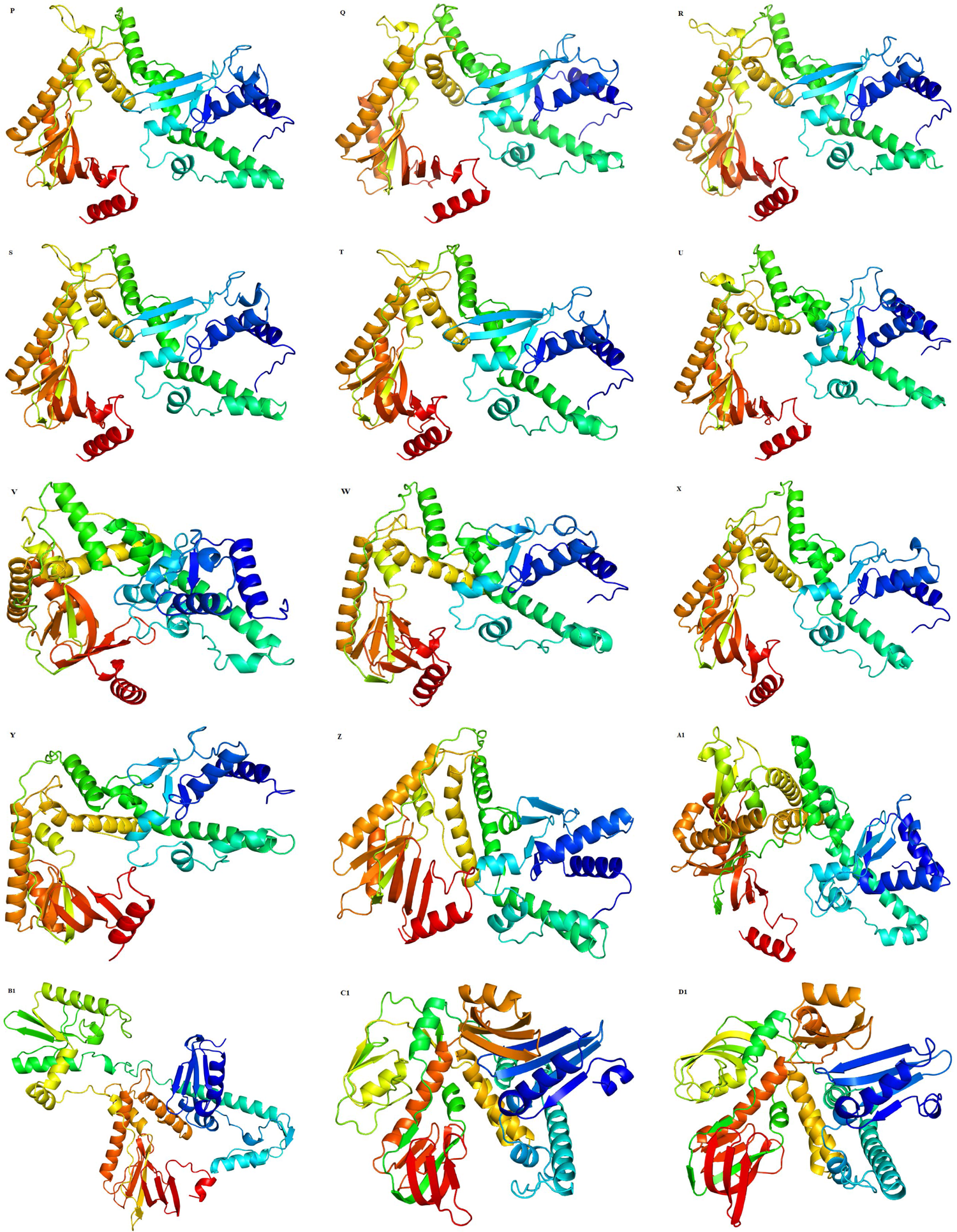
Dendrogram of the δ catalytic subunits and Family B polymerases of NCLDV. The dendrogram is based on the pairwise Z score comparisons and calculated using DALI. GenBank accessions are specified for each predicted protein structure. Protein structure prediction was done with ColabFold using MMseqs2 (47, 48). The length of the predicted structures ranged from 400 to 421 amino acids. Only unrelaxed rank 1 predicted protein structures were used to calculate the dendrogram. The dendrogram is rooted between the viral and cellular clades according to the topology of Fig. 1, as well as in published data (22, 39).

### The morphogenetic module and “synapomorphic” proteins

Phylogenetic relationships of the FtsK-HerA superfamily ATPases are complex and incompletely resolved. Analysis of the presence and absence of characteristic secondary structure elements beyond the conserved core consisting of a seven-stranded β-sheet, as well as synapomorphic amino acid substitutions, made it possible to delineate a clade consisting of the HerA, VirB3, VirB4, VirD4, FtsK, and A32-like (NCLDV) ATPases (52, 53). We used the anchor amino acid substitutions identified by Iyer and colleagues (52) to create an alignment of the highly conserved Helix-1 - Strand 4 (sGsGKo – Q) fragment. The phylogeny of the FtsK-HerA superfamily clades coincided with the Iyer and colleagues’ results (52) (Fig. S47, Align. S51, Data S58). This allowed us to reconstruct a tree consisting of the cellular and A32-like homologs (Fig. 6, Align. S52, Data S59). The obtained topology showed that the FtsK and A32-like ATPases formed a clade with high statistical support. *Phycodnaviridae* s.l. and the rest of the NCLDV families formed sister clades (Table 1). We also constructed a tree without cellular orthologues (Fig. S48, Align. S53, Data S60). The resulting topologies of both trees showed that the MAPIM and AP clades were sisters whereas MAPIMAP formed a sister clade to *Mimiviridae* s.l. These findings corresponded to the topologies of the eight proteins described above (Table 1). A similar direction of evolution demonstrated the FtsK-HerA superfamily ATPases dendrogram (Fig. S49, Data S61). However, there were two differences between the topologies of the dendrogram and phylogenic trees. Firstly, on the dendrogram, the *Emiliania huxleyi virus 86* prediction was localized basally to all crown NCLDVs, and the *Pandoraviridae* + *Prasinovirus* clade formed a sister clade to all other NCLDV families whereas on the phylogenetic tree the phycodnaviruses s.l. retained their monophyly (Fig. 6). Secondly, on the dendrogram, the AP clade was sister to the *Mimiviridae* s.l. clade, while on the phylogenetic trees the AP clade was sister to the MAPIM clade (Fig. 6, Fig. S48). Next, we showed the phylogenies of the DJR MCP (Fig. S50, Align. S54, Data S62) and VLTF-2 (Fig. S51, Align. S55, Data S63) proteins. Both phylogenies were constructed without using of outgroup and for this reason, both trees were rooted according to the direction of evolution of individual proteins presented above. Notably, the topologies of both “synapomorphic” proteins showed sister relationships between the AP and MAPIM clades, as well as between the joint AP + MAPIM and *Mimiviridae* clades. However, the absence of outgroups that determined the direction of evolution of NCLDV made it possible to reroot both trees according to the Koonin and Yutin hypothesis (10).

**Fig. 6.**
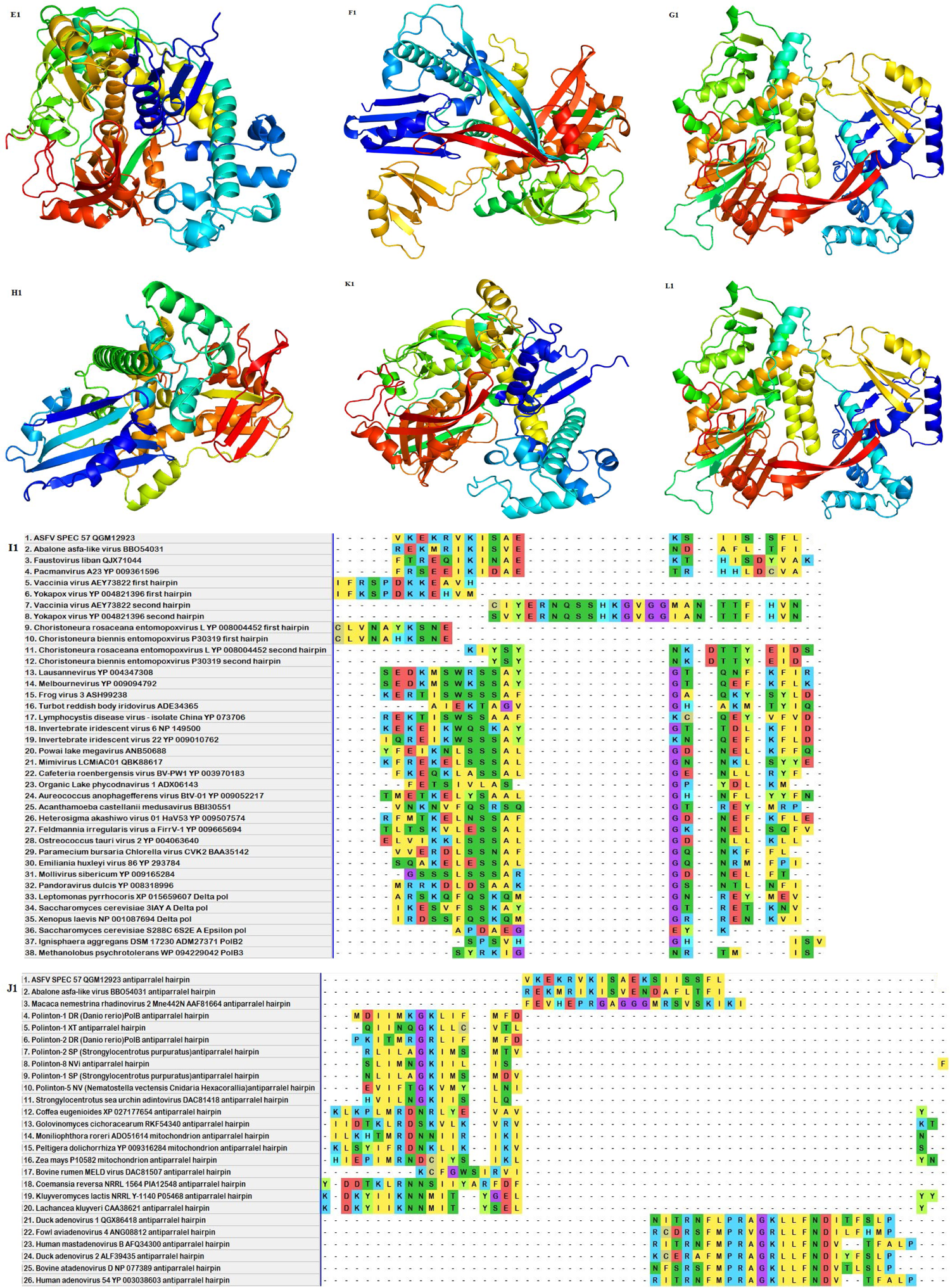
Maximum likelihood phylogenetic tree of the PilT, FtsK, HerA, VirB3, VirB4, and A32-like ATPases. The tree was constructed using the IQ TREE program (version 1.6.12) under the LG+R3 model and rooted according to the Fig. S47 topology, as well as in published data (50). The corresponding alignment of the Helix-1 – Strand 4 (sGsGKo – Q) fragment was built using the MUSCLE algorithm. Statistical support of the partitions, measured by SH-aLRT branch tests (1,000 replicates), approximate Bayes, and ultrafast bootstrap (1,000 replicates), is shown at the nodes. Scale bars represent the number of amino acid substitutions per site.

Finally, we showed the N and C terminal jelly roll barrels (tjrb) predictions dendrograms (Figs. S52, S53, Data S64, S65), and the dendrogram of the full-size DJR MCP predictions (Fig. S54, Data S66) whose tertiary structures were predicted using ColabFold, as well as the dendrogram of the full-size DJR MCP predictions (Fig. 7, Data S67) obtained using I-TASSER (54–56) and combined with the NCLDV crystallographic structures deposited in the Protein Data Bank (57). All predictions were calculated without removing any amino acid residues. Unlike most of the dendrograms presented above, and, conversely, according to most individual phylogenies, the topologies of the N and C tjrbs showed sister localization of the MAPIM and AP clades. However, in contrast to both N and C tjrbs dendrograms (Figs S52, S53), the topology of the full-size DJR MCPs dendrogram (Fig. S54) placed the AP clade sisterly to the joint MAPIM + *Mimiviridae* s.l. clade. In its turn, in contrast, the dendrogram S54 topology shown in Figure 7 placed the AP and MAPIM clades sisterly, thereby demonstrating topological congruence with the topologies of Figures S52 and S53. We excluded the ColabFold predictions from the calculation of the dendrogram corresponding to Figure 7, since the *Mimiviridae* s.l. and MAPIM predictions obtained using ColabFold demonstrated attraction to each other, separating from the rest of the members of their clades (Data S68).

**Fig. 7.**
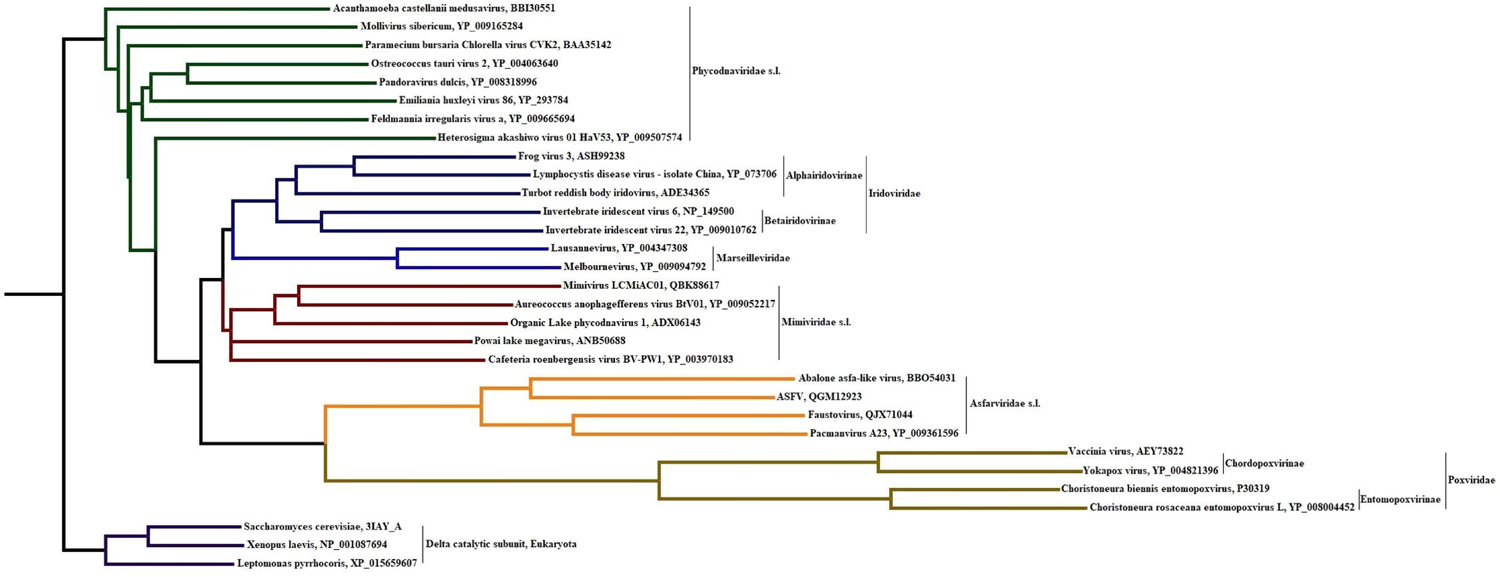
Dendrogram of the NCLDV DJR MCPs. The dendrogram is based on the pairwise Z score comparisons calculated using DALI. Protein structure prediction was done with I-TASSER (54–56). The dendrogram was rooted between *Phycodnaviridae* and MAPIMAPM clades according to the topologies of all phylogenetic trees with outgroups presented in this work. GenBank accessions are given for the sequences used for the I-TASSER predictions, as well as for the crystallographic structures deposited in the PDB. The I-TASSER prediction was carried out without removing any amino acid residue within the sequences.

Two important generalizations followed from the analysis of 18 individual proteins. Firstly, in 11 cases (D5-like primase, DNA topoisomerase II, PolB, A32-like ATPase, Metallopeptidase WLM, mRNA capping enzyme, PCNA, RNAP α, RNAP β, RuvC, and TFIIB) where outgroups were used, genes were encoded by phycodnaviruses s.l. and were not transferred secondary, between *Phycodnaviridae* s.l. and a clade consisting of other NCLDVs statistically well-supported sister relationships were found (Table 1). Secondly, the sister relationships between the AP and MAPIM clades, as well as between the MAPIMAP and *Mimiviridae* s.l. clades were observed in 12 cases (Table 1).

Dendrograms calculated using outgroups showed an evolutionary trajectory similar to individual phylogenies. On dendrograms phycodnaviruses s.l. either formed a sister clade to other NCLDVs or were localized basally to crown NCLDVs. The topological position of the AP clade was a greater variable compared to individual phylogenies (see above).

### The branching order of NCLDV. The SPR superdendrogram and SPR supertree supported the sister relationship between *Phycodnaviridae* s.l. and other NCLDVs and clarified the position of the *Asfarviridae*/*Poxviridae* clade

One of the approaches to solving topological disagreements arising from phylogenetic analyses of various genes and proteins is the construction of supertrees. Superdendrograms can be constructed for the same reason. In our case, the topological conflict referred to the position of the AP clade.

To construct a subtree prune-and-regraft distance (SPR) superdendrogram (58), we generated 12 dendrograms of 9 core proteins using ColabFold (Data S69-S80). The topology of the superdendrogram showed sister relationships between the AP and MAPIM clades, as well as between the joint MAPIMAP and *Mimiviridae* s.l. clades (Fig. 8, Data S81). However, monophyletic *Phycodnaviridae* s.l. were split into two clades. The first clade consisted of the *Phaeovirus*, *Coccolithovirus*, and *Pandoraviridae* predictions and was localized sisterly to the joint MAPIMAPM clade. The second clade consisted of the *Raphidovirus*, *Acanthamoeba castellanii medusavirus*, *Chlorovirus*, *Prasinovirus*, and *Mollivirus sibericum* predictions and was localized basally to the crown NСLDV families.

**Fig. 8.**
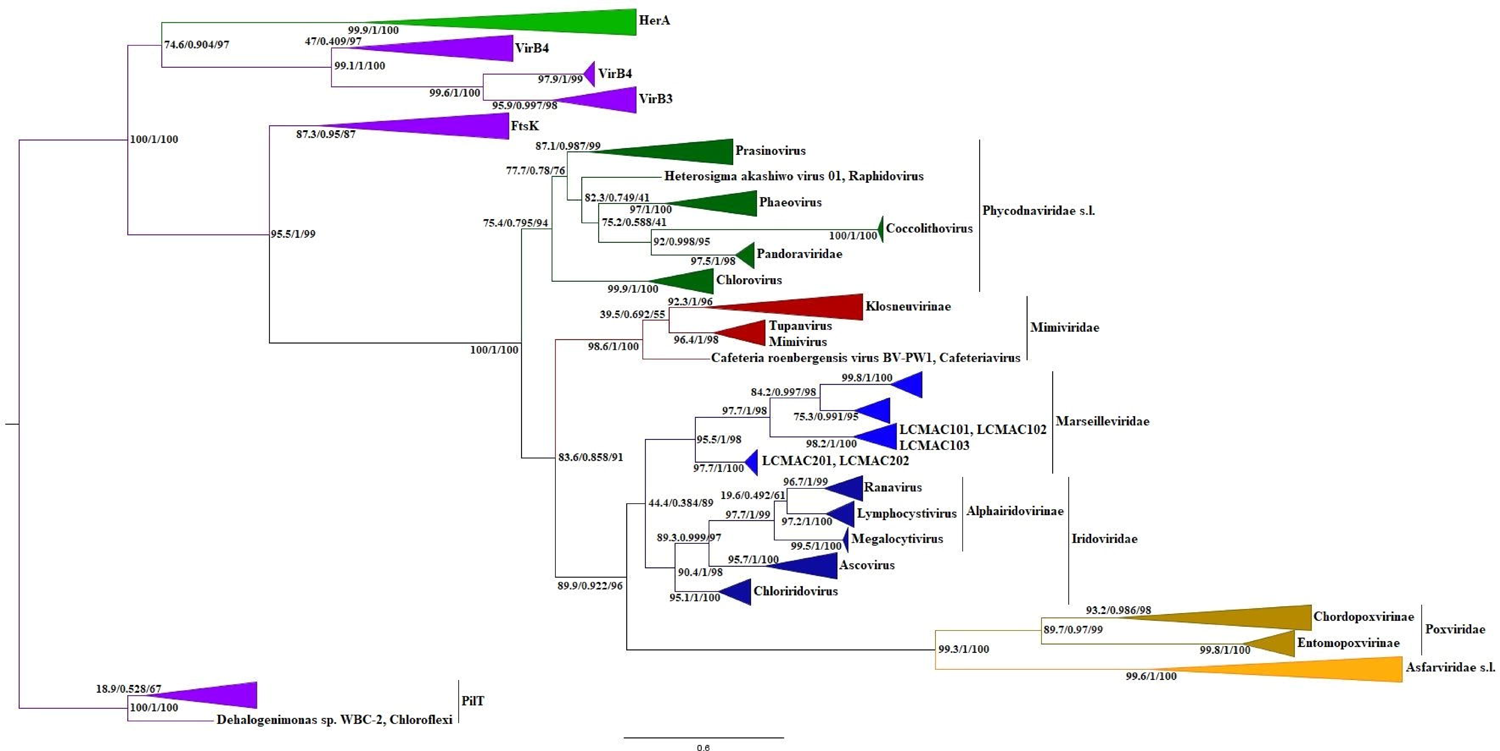
SPR superdendrogram. The dendrogram is based on 12 ColabFold predictions derived from the 9 NCLDV core proteins (PolB, RNAP α, RNAP β, PCNA, Topoisomerase II, TFIIB, A32-like ATPase, Uracil DNA glycosylase, as well as N and C terminal jelly roll barrels of the DJR MCPs). The superdendrogram was rooted between *Eukaryota* and NCLDV according to the topologies of all individual phylogenies containing the cellular and NCLDV clades presented in this work. The input dendrograms of the PolB, RNAP α, RNAP β, PCNA, Topoisomerase II, and TFIIB predictions contained the corresponding eukaryotic outgroups. The remaining input dendrograms were calculated without using outgroups.

To construct the SPR supertree, we generated the 23 maximum likelihood trees of the 13 core proteins with a limited (1-4 sequences from each genus or subfamily) taxonomic composition, using the standard bootstrap method for statistical support of nodes (Align. S56-S78, Data S82-S104). In contrast to the superdendrogram, phycodnaviruses s.l. retained their monophyly and were localized sisterly to the joint MAPIMAPM clade (Fig. 9, Data S105). Meanwhile, the topological relationships between the AP and MAPIM, as well as between the MAPIMAP and *Mimiviridae* s.l. clades repeated the topology of the superdendrogram, as well as of the vast majority of individual phylogenies (see above).

**Fig. 9.**
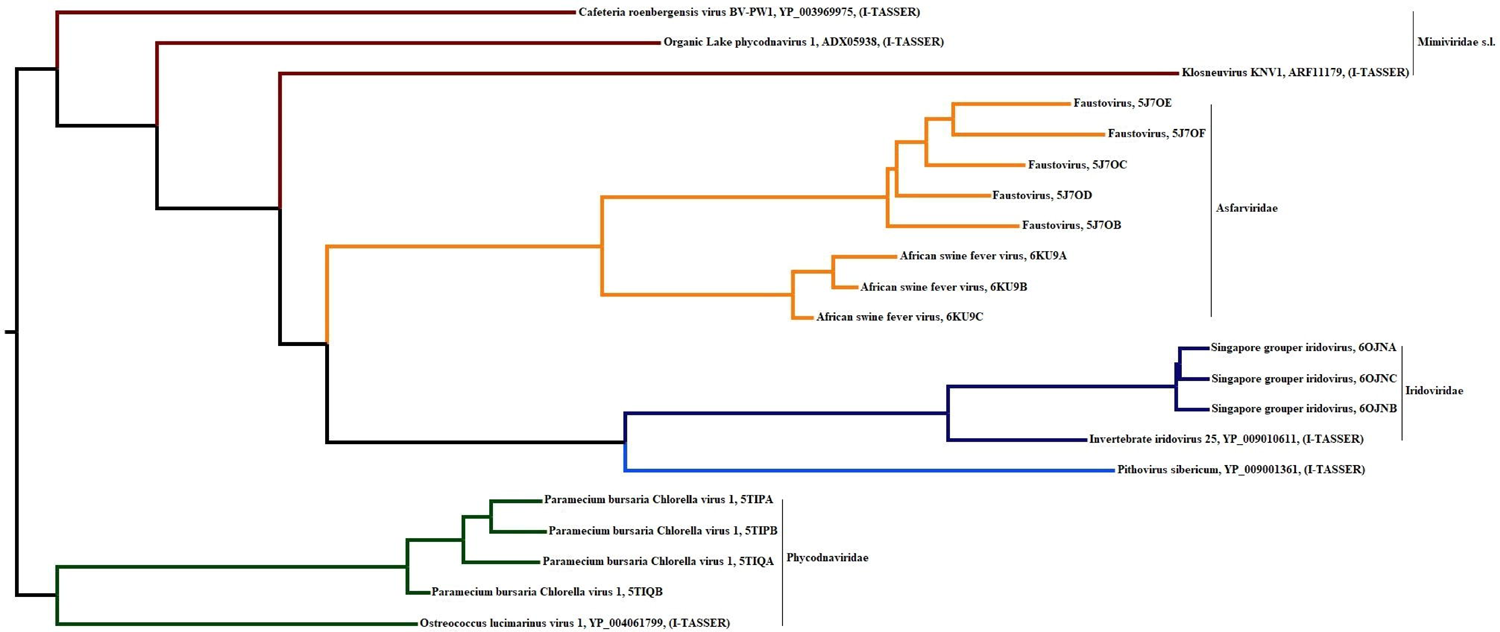
SPR supertree. The supertree was constructed based on the 23 maximum likelihood (ML) trees of the 13 core proteins (A32-like ATPase, AP (apurinic) endonuclease family 2, D5-like primase, mRNA capping enzyme, PolB, RNAP α, RNAP β, DJR MCP, SFIII helicase, TFIIB, Uracil DNA glycosylase, VETF, and VLTF-2). The supertree was constructed with a bootstrap threshold equal to 80 and rooted between *Eukaryota* and NCLDV according to the topologies of all individual phylogenies containing the cellular and NCLDV clades presented in this work. The input trees of the D5-like primase, mRNA capping enzyme, PolB, RNAP α, RNAP β, and TFIIB were constructed using the corresponding eukaryotic outgroups. The remaining input trees were constructed without using outgroups.

### The branching order of NCLDV. The concatenated phylogeny supported the sister relationship between *Phycodnaviridae* s.l. and other NCLDVs and clarified the position of the *Asfarviridae*/*Poxviridae* clade

Concatenation is an alternative way to solve topological disagreements among individual phylogenies. Alignments of the a) 16 individual proteins of eukaryotic/protoeukaryotic, bacterial, and unknown origin with and without using of eukaryotic outgroups (Table S4, Supermatrixes 1 and 2), b) 6 individual proteins of eukaryotic/protoeukaryotic origin with and without using of eukaryotic outgroups (Table S4, Supermatrixes 3 and 4), c) 4 individual proteins of bacterial origin with and without using of bacterial outgroups (Table S4, Supermatrixes 5 and 6), and d) 3 individual proteins of unknown origin without outgroups (Table S4, Supermatrix 7) were built for concatenation. Thus, based on these seven concatenated alignments, we constructed seven different concatenated phylogenies (Fig. 10, Figs. S55-S60, Aligns. S79-S85, Data S106-S112).

**Fig. 10.**
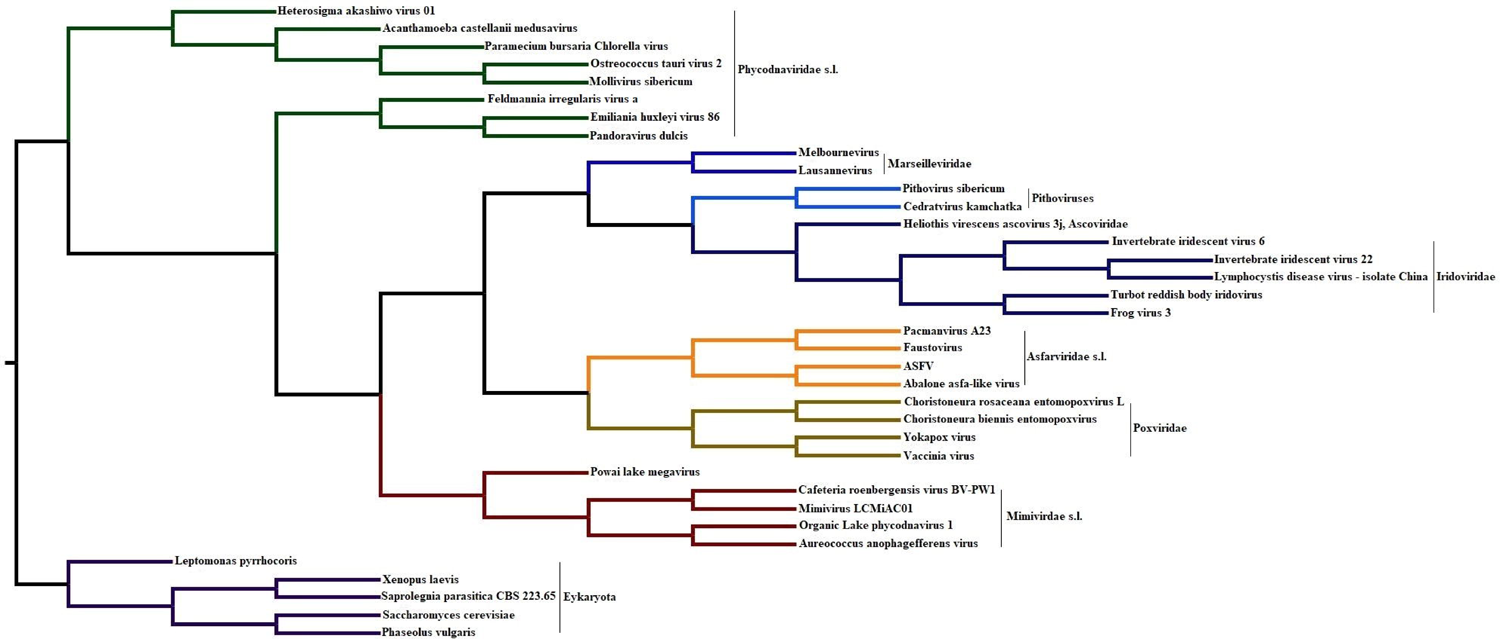
Maximum likelihood phylogenetic tree of the NCLDVs. The tree was constructed via concatenation of the NCLDV 16 core protein (D5-like primase, PolB, mRNA capping enzyme (large subunit), PCNA, RNAP α, RNAP β, TFIIB, Topoisomerase II, FtsK-HerA superfamily ATPase, AP (apurinic) endonuclease family 2, Holliday junction resolvase, Uracil-DNA glycosylase, SFIII helicase, DJR MCP, VETF, and VLTF-2) alignments consisting of NCLDV and eukaryotic homologs using the IQ TREE program (version 1.6.12) under the EX_EHO+F+I+R4 model. The tree was rooted between *Eukaryota* and NCLDV according to the topologies of all individual phylogenies with outgroups presented in this work. The supermatrix was built via head-to-tail linking of already aligned (MUSCLE) individual protein alignments. Statistical support of the partitions, measured by SH-aLRT branch tests (1,000 replicates), approximate Bayes, and ultrafast bootstrap (1,000 replicates), is shown at the nodes. Scale bars represent the number of amino acid substitutions per site.

The use of eukaryotic and bacterial outgroups showed that the direction of evolution of the mixed, eukaryotic, bacterial, and “synapomorphic” concatenated trees repeated the branching order of individual phylogenies and dendrograms, as well as supertree and superdendrogram. That is, the *Phycodnaviridae* s.l. and MAPIMAPM clades showed a sister relationship. Moreover, all seven concatenated phylogenies repeated the topological relationships between the *Mimiviridae* s.l., MAPIM, and AP clades, i.e. the AP clade with high statistical support was localized sisterly to the MAPIM clade, and the MAPIMAP clade, in turn, with high statistical support, was localized sisterly to the *Mimiviridae* s.l. clade.

### Phylogenetic relationships among proteins (genes) of NCLDV, *Preplasmiviricota*, several classified and unclassified viruses, and MGEs

#### The replication module

Our study showed that the Exo I–MotifC fragments belonging to *Adenoviridae*, *Adintoviridae*, *Bidnaviridae*, *Parvoviridae*, CLPs, MLPs, and Polintons formed a clade and localized sisterly to pol δ (Fig. S61, Align. S86, Data S113). Similarly, the Exo I–MotifC fragments of *Herpesviridae* formed a sister clade to pol δ (Fig. S62, Align. S87, Data S114). However, the Exo I–MotifC fragments of *Hytrosaviridae* were found within the pol δ clade, suggesting their origin from within the eukaryotic δ catalytic subunits (Fig. S63, Align. S88, Data S115). Thus, our results for herpesviruses and hytrosaviruses were consistent with previously published data (20–22). We removed the pol δ sequences from the alignments S86 and S87, assuming that if the topological positions of viral clades were not the results of homoplasies, then the viral clades will retain their previous topological positions. Indeed, the *Herpesviridae* and *Adenoviridae*/Polintons clades retained their topological positions, thus confirming their evolutionary relationship with the eukaryotic δ catalytic subunits (Figs. S64-S65, Aligns. S89-S90, Data S116-S117).

The sister relationships between the *Herpesviridae*, *Adenoviridae*/Polintons, and NCLDV clades on the one hand and the eukaryotic δ catalytic subunits on the other, led to the need to check the phylogenetic relationships among the *Herpesviridae*, *Adenoviridae*/Polintons, and NCLDV Exo I– MotifC fragments. Reconstruction of a phylogenetic tree, consisting of the pol δ, NCLDV, and *Herpesviridae* Exo I–MotifC fragments, showed that *Herpesviridae* formed a sister clade to the joint *Mimiviridae* s.l. + MAPIM clade with high statistical support (Fig. S66, Align. S91, Data S118). It is important to note that the topological position of herpesviruses (Fig. S66) coincided with the topological position of seven metagenomic sequences (Fig. 2) which we identified as sequences related to *Herpesviridae* and pol δ. These findings suggested that the ancestral Exo I–MotifC fragment of *Herpesviridae* presumably was acquired from the lineage leading to the last common ancestral Exo I–MotifC fragment of the *Mimiviridae* s.l., MAPIM, and AP clades.

Next, we showed that the Exo I–MotifC fragments of *Adenoviridae*, *Adintoviridae*, *Bovine rumen MELD virus*, *Sheep rumen MELD virus*, *Bidnaviridae*, *Parvoviridae*, CLPs, MLPs, and Polintons formed a clade with the highest statistical support, whose last common ancestral fragment presumably originated from within NCLDV (Fig. 11, Fig. S67, Alig. S92-S93, Data S119-S120). These results were obtained using both the MUSCLE and MAFFT algorithms. However, on both trees, the statistical support of the joint *Mimiviridae* s.l. + *Adenoviridae*/Polintons clade was low. We added to the analysis the Exo I–MotifC fragments of *Monoraphidium MELD virus SAG 48.87*, *Rhizophagus glomeromycete fungus MELD virus 6100*, *Blastocystis MELD virus 3996*, and *MELD virus sp. ctWXX4* and reconstructed the tree (Fig. S68, Align. S94, Data S121). During the building of this alignment, we endeavored to maximize the preservation of conservative vertical blocks and amino acid substitutions of the DALI structure-based sequence alignment, highlighted in color in Supplementary Figure S1 (22). The *Adenoviridae*/Polintons clade was localized within the MAPIM clade. (Fig. S68, Align. S94, Data S121). Thus, the resulting topology contrasted with the previous two trees. We used as a template the alignment S10 based on the mTM-align multiple protein structure alignment (Align.S8, see above) to align the Exo I–MotifC fragment of NCLDVs and metagenomic sequences (43) with the prealigned within *Adenoviridae*/Polintons lineage framework homologous sequences of *Adenoviridae*, *Adintoviridae*, CLPs, MLPs and Polintons using the MUSCLE v5 software. The resulting tree showed the origin of the Exo I - MotifC fragments of the *Adenoviridae*/Polintons clade from within NCLDV but localized them within *Mimiviridae* s.l. (Fig. 12, Align. S95, Data S122).

**Fig. 11.**
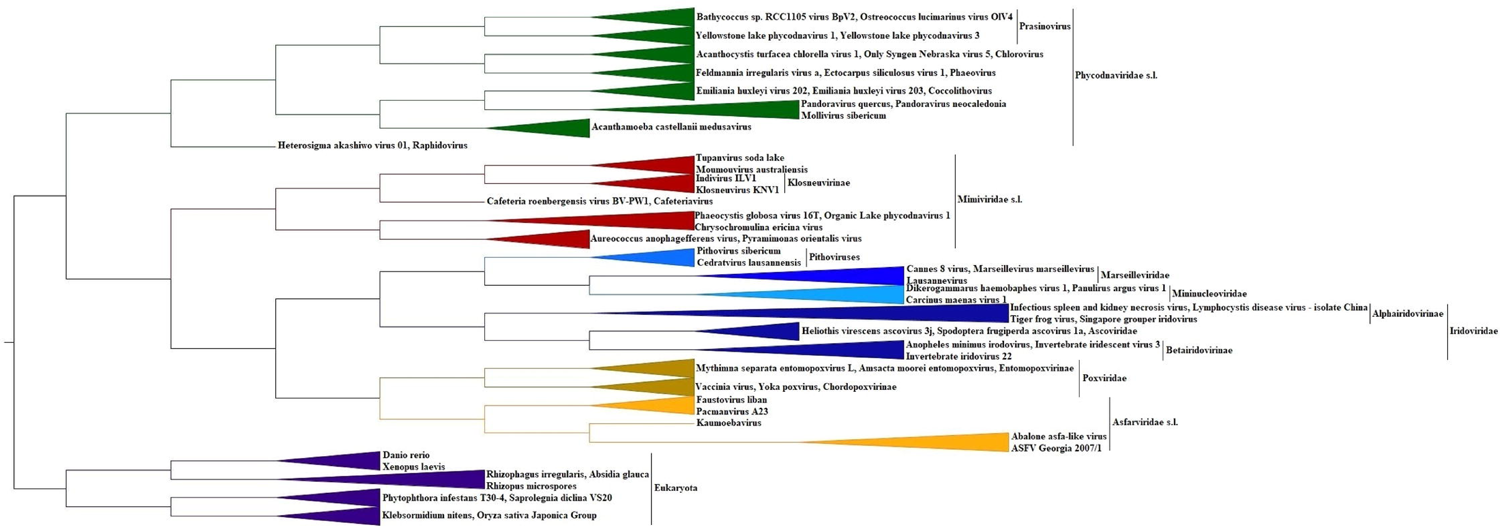
Maximum likelihood phylogenetic tree of Family B DNA polymerases. The tree consists of the eukaryotic δ catalytic subunits, the NCLDV, *Adenoviridae*, *Adintoviridae*, *Bidnaviridae*, *Parvoviridae*, *Bovine rumen MELD virus*, *Sheep rumen MELD virus*, Polintons, Mitochondrial linear plasmids, and Cytoplasmic linear plasmids Family B DNA polymerases. The tree was constructed using the IQ TREE program (version 1.6.12) under the LG+F+I+R4 model and rooted according to Fig. 1, Fig. 2, and Figs. S9-S10 topologies. The corresponding alignment was built using the MUSCLE algorithm. Statistical support of the partitions, measured by SH-aLRT branch tests (1,000 replicates), approximate Bayes, and ultrafast bootstrap (1,000 replicates), is shown at the nodes. Scale bars represent the number of amino acid substitutions per site.

**Fig. 12.**
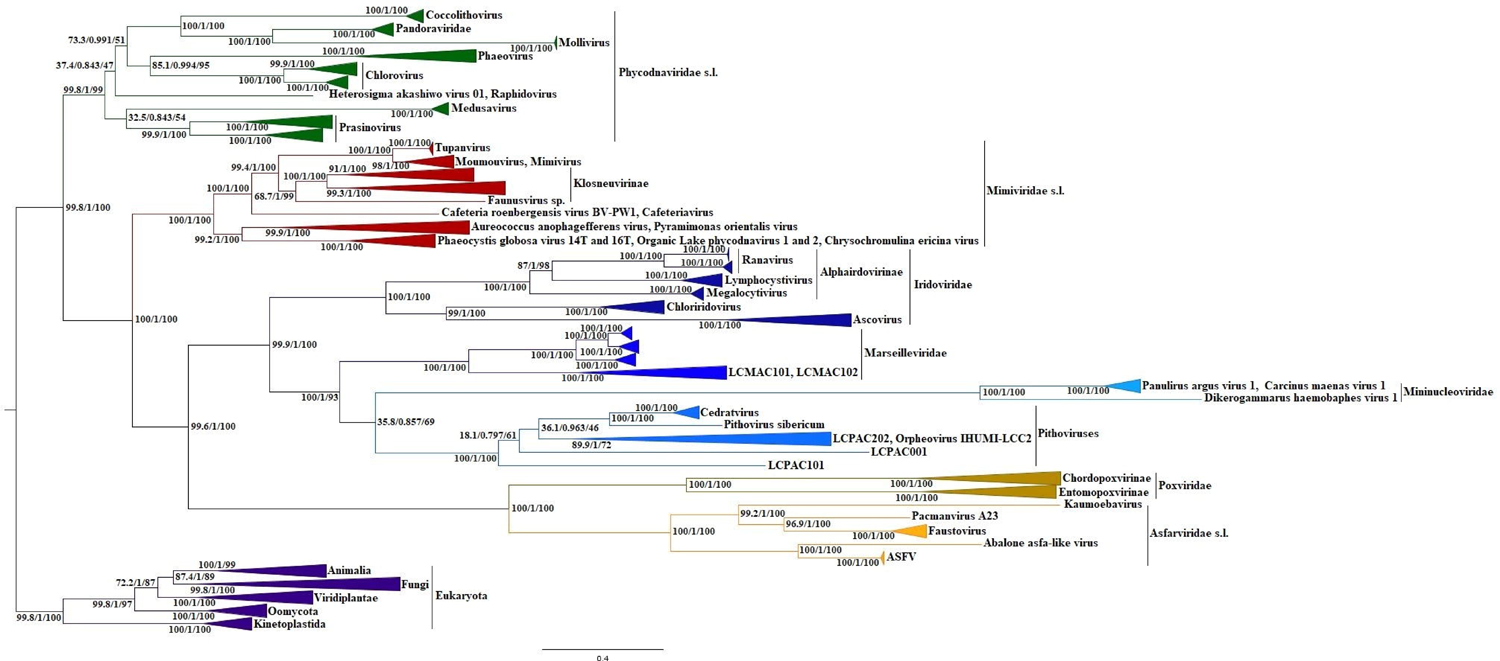
Maximum likelihood phylogenetic tree of the eukaryotic δ catalytic subunits, NCLDV, viral metagenomic, and *Adenoviridae*/Polintons clade Family B DNA polymerases. The tree was constructed using the IQ TREE program (version 1.6.12) under the LG+F+I+R4 model and rooted according to the topologies of Fig. S1, Fig. 1, and Fig. 2, as well as in published data (22, 39). The corresponding alignment of the Exo I – MotifC fragment was built using the MUSCLE v5 software under the permutation none, perturbation default parameters, and using the alignment S10 as a template. Square brackets indicate the number of metagenomic sequences belonging to a given clade. Statistical support of the partitions, measured by SH-aLRT branch tests (1,000 replicates) and approximate Bayes, is shown at the nodes. Scale bars represent the number of amino acid substitutions per site.

We removed the metagenomic sequences of the Exo I–MotifC fragment and a) the identified sequences of *Mimiviridae* s.l., b) the identified sequences of the *Mimiviridae* s.l. and AP clades from the alignment S95 and reconstructed two trees. On both trees, the *Adenoviridae*/Polintons clade was localized sisterly to mininucleoviruses: a) without mimiviruses (Aligns. S96, Data S123), and b) without mimiviruses and the AP clade (Align. S97, Data S124).

We predicted, visualized, and explored the morphology of the Exo I - MotifC fragment of *Macaca nemestrina rhadinovirus 2* (*Gammaherpesvirinae*), *Adenoviridae*, *Adintoviridae*, CLPs, MLPs, and Polintons PolBs (Fig. 4A1-4H1, Table S2). The predictions of *Macaca nemestrina rhadinovirus 2*, adenoviruses, CLPs, MLPs, *Bovine rumen MELD virus*, *Strongylocentrotus sea urchin adintovirus*, and Polintons showed the presence of a beta-hairpin typical for the NCLDV, pol δ, pol ε, PolB2, and PolB3 predictions (Fig. 4A1-4H1, J1, Table S2, Align. S98). However, closely related to Polintons adintoviruses and MELD viruses (59) were separated according to this morphological criterion. In particular, in *Sheep rumen MELD virus* and adintoviruses excepting *Strongylocentrotus sea urchin adintovirus*, the beta-hairpin was missing (Fig. 4 K1-4L1, Table S2). We did not rule out the possibility of a trend toward the loss of a beta-hairpin in this group of viruses and mobile elements. The length and position of beta-hairpins of the viruses and MGEs under study relative to the NCLDV beta-hairpins were consistent with the phylogenetic position of their carriers relative to NCLDV (Fig 4 J1). The beta-hairpin of *Macaca nemestrina rhadinovirus 2* was about the same length as hairpins of NCLDV and located in the β16-β17 fragment, which was consistent with the assumption of the origin of *Herpesviridae* polymerases from within NСLDV (Fig. 4J1). Among the sequences of the *Adenoviridae*/Polintons clade, only in adenoviruses the N termini of the beta-hairpins overlapped with the C termini of the beta-hairpins of NCLDV and pol δ (Fig 4 J1, Align. S96). This observation was in agreement with the basal position of *Adenoviridae* to CLP, MLP, and Polintons (Figs. 11-12, Figs. S67-68). According to the alignment S98 the beta-hairpins of MLPs, CLPs, and Polintons were shifted towards the N terminus of the Exo I – MotifC fragment and did not overlap with the beta hairpins of NCLDV and pol δ.

Next, we showed two dendrograms indicating the origin of the Exo I - MotifC fragment of the *Adenoviridae*/Polintons clade from within NCLDV. The dendrogram calculated without using the *Asfarviridae* s.l. predictions localized the *Adenoviridae*/Polintons clade sisterly to the joint *Mimiviridae* s.l. + MAPIM clade (Fig. 13, Data S125); while the dendrogram calculated using the predictions of *Asfarviridae* s.l. localized the *Adenoviridae*/Polintons clade sisterly to asfarviruses (Data S126).

**Fig. 13.**
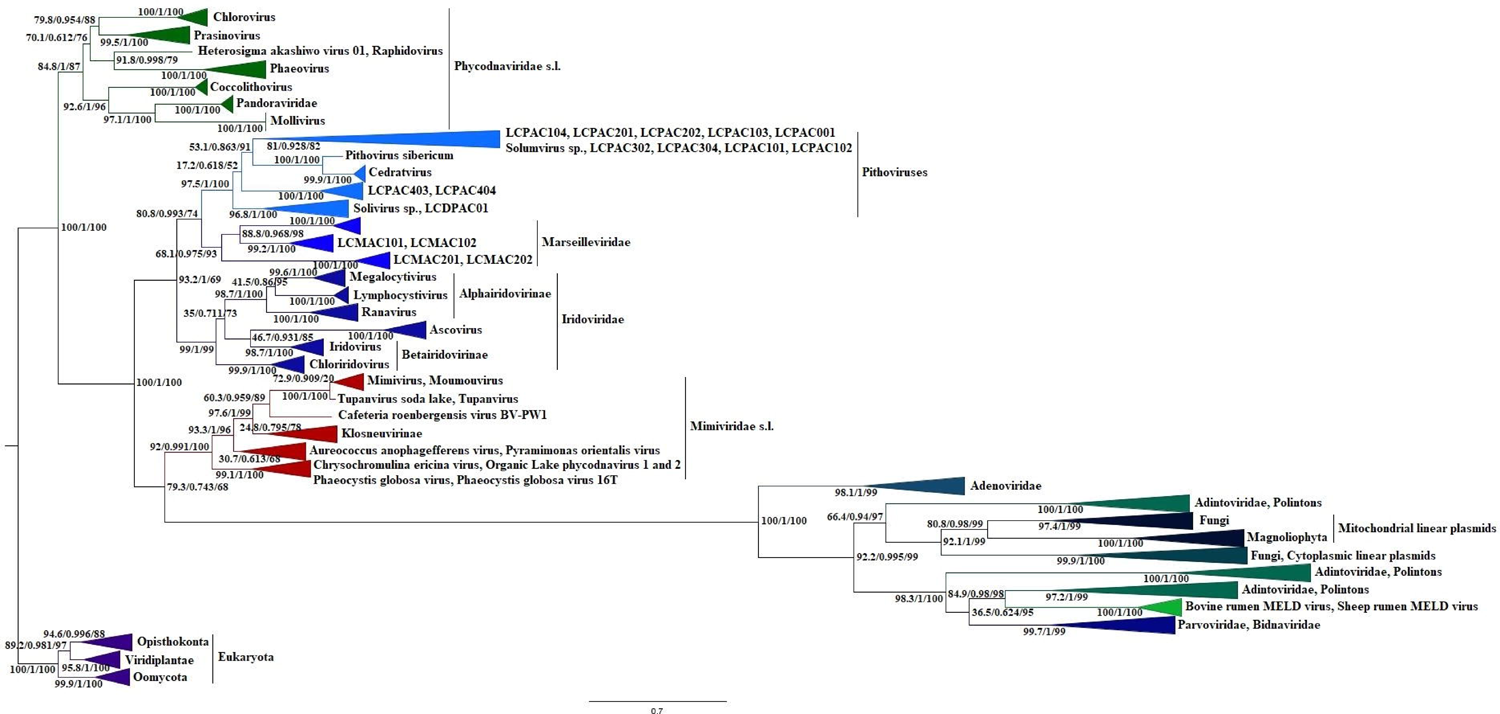
Dendrogram of Family B DNA polymerases. The dendrogram consists of the eukaryotic δ catalytic subunit, NCLDV, *Adenoviridae*, *Adintoviridae*, *Bovine rumen MELD virus*, *Sheep rumen MELD virus*, Polintons, MLP, and CLP Family B DNA polymerase predictions and based on the pairwise Z score comparisons calculated using DALI. GenBank accessions are specified for each predicted protein structure. Protein structure prediction was done with ColabFold using MMseqs2 (47, 48). The length of the predicted structures ranged from 400 to 615 amino acids. Only unrelaxed rank 1 predicted protein structures were used to calculate the dendrogram. The dendrogram was rooted between the viral and cellular clades according to the Fig 1 and 2 topologies.

Thus, we have little phylogenetic statistical arguments for the affinity of the Exo I - MotifC fragment of the *Adenoviridae*/Polintons clade with mimiviruses s.l. However, all obtained phylogenies and dendrograms confidently indicate the origin of the Exo I - MotifC fragment of the *Adenoviridae*/Polintons clade from within NCLDV.

Finally, we compared the topological position of the Exo I–MotifC fragment of the *Adenoviridae*/Polintons clade with the position of the same fragment of the prokaryotic *Preplasmiviricota*. The Exo I – MotifC fragment of the prokaryotic *Preplasmiviricota*, *Ampullaviridae*, *Fuselloviridae*, *Pleolipoviridae*, *and Salterprovirus* along with the homologous of *Euryarchaeota*, *Thaumarchaeota*, *and Firmicutes* formed a clade with high statistical support, which was localized sisterly to pol ε (Fig. S69, Alig. S99, Data S127). Thus, the Exo I - MotifC fragment of prokaryotic *Preplasmiviricota* has an origin unrelated to pol δ and NСLDV. In other words, according to the Exo I – MotifC fragment phylogenies *Preplasmiviricota* are polyphyletic.

D5-like primases of *Sputnikvirus* (*Lavidaviridae*, *Preplasmiviricota*) formed a sister clade to the *Iridoviridae*/*Ascovirus* clade (Fig. S70, Align. S100, Data S128). Removal of the *Iridoviridae*/*Ascovirus* sequences from the alignment did not change the topological localization of sputnikviruses (Fig. S71, Align. S101, Data S129). Thus, we had reason to suspect that the D5-like primases of sputnikviruses originated from within the joint *Mimiviridae* s.l. + MAPIM clade, similar to the Exo I – MotifC fragment of PolBs.

Tree, consisting of the NCLDV D6/D11 (SFII) helicases, the DEAD-like helicases of *Fungi* (CLPs) and *Physcomitrella patens* (*Bryophyta*, chromosomes 3 and 13) showed that the fungal and *P. patens* sequences were localized within the MAPIM clade (Fig. 14, Align. S102, Data S130). The removal of the sequences of *Iridoviridae* and pithoviruses from the alignment S102, taking into account the topology of Fig. 3, that is, in fact, the reciprocal removal of sister clades, showed that the topological localization of the joint CLP + *P. patens* clade did not change, which spoke in favor of the absence of homoplasies (Fig. S72, Align. S103, Data S131). The topological position of the *P. patens* chromosomal DEAD-like helicases suggests their origin from CLPs. Along with PolB and D5-like primase, the DEAD-like helicases of CLPs were the third protein (gene) of the *Adenoviridae*/Polintons clade, presumably have originated from within NCLDV.

**Fig. 14.**
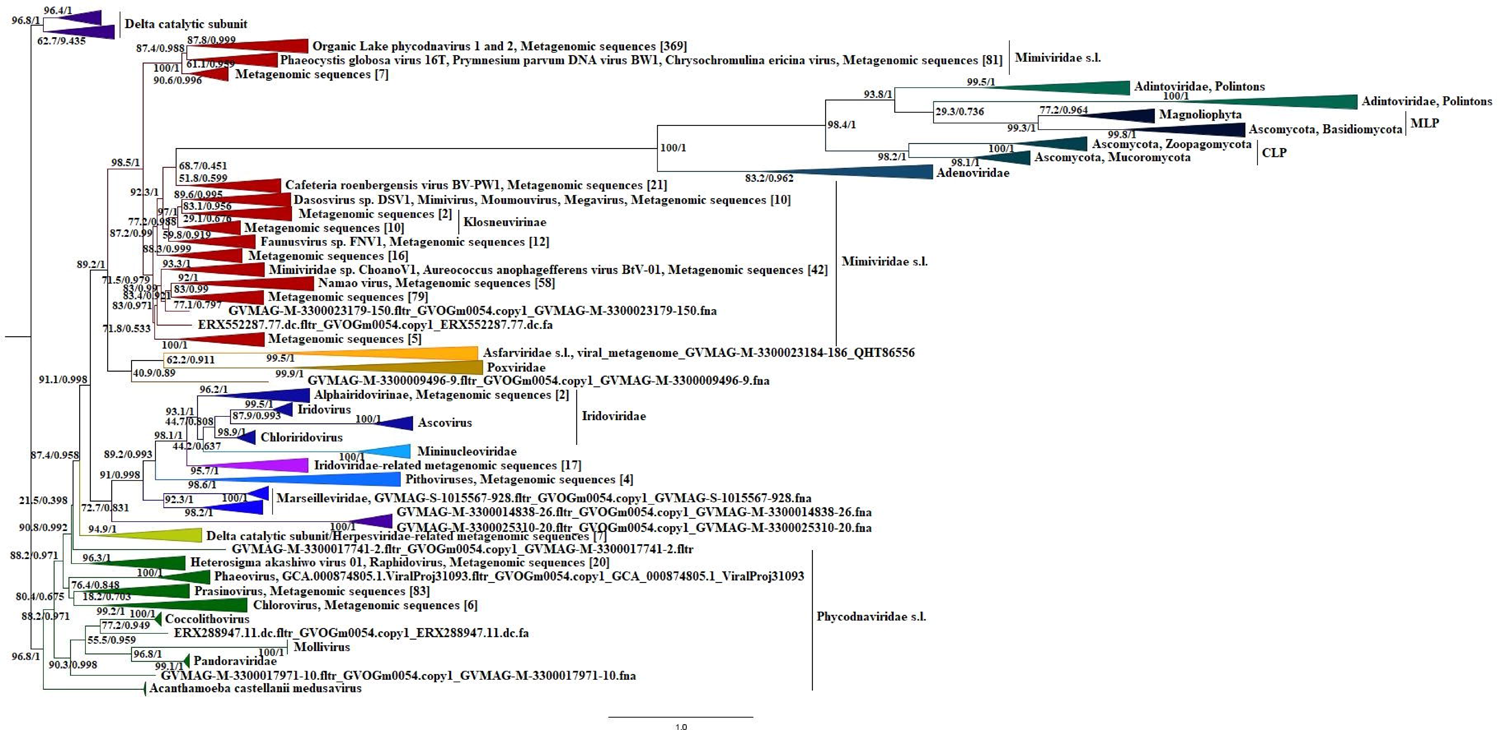
Maximum likelihood phylogenetic tree of the D6/D11 and DEAD-like helicases. The tree consists of the *Chlorovirus*, *Mimiviridae* s.l. *Marseilleviridae*, *Cedratvirus*, *Pithovirus sibericum*, *Iridoviridae*/*Ascoviridae*, *Asfarviridae* s.l., *Poxviridae*, CLP, and *Physcomitrella patens* (chromosomes 3 and 13) protein sequences. The tree was constructed using the IQ TREE program (version 1.6.12) under the LG+C20+F+I+R4 model and rooted between chloroviruses and the clade consisting of the rest of D6/D11 + CLP + *P. patens* homologs. The corresponding alignment was built using the MUSCLE algorithm. Statistical support of the partitions, measured by SH-aLRT branch tests (1,000 replicates), approximate Bayes, and ultrafast bootstrap (1,000 replicates), is shown at the nodes. Scale bars represent the number of amino acid substitutions per site.

The topology of the phylogenetic tree, consisting of the F1055L (SFII helicases) homologs of ASFV of *Mimiviridae*, *Marseilleviridae*, *Asfarviridae*, as well as *Herpesvirales*, localized *Herpesvirales* sisterly to *Asfarviridae* (Fig. S73, Align. S104, Data S132). However, the statistical support of the node combining *Asfarviridae* and *Herpesvirales* was low. Moreover, the *Herpesvirales* branch was long and could contain homoplasies with the asfarviruses branch, which could lead to an artificial topology. We removed the sequences of asfarviruses from alignment S104 and reconstructed the tree (Fig. S74, Align. S105, Data S133). As can be seen from Fig. S74, the topological position of *Herpesvirales* remained unchanged. The constancy of the topological position of herpesviruses spoke against the presence of potential homoplasies, as well as suggested the origin of the *Herpesvirales* homologs from within NCLDV, as in the case of *Herpesviridae* polymerases.

Next, we included the *Herpesviridae* and *Lavidaviridae* SFIII helicases in our analysis. The result suggested that the LCA genes of *Lavidaviridae* and *Herpesviridae* originated from within NCLDV and localized sisterly to the MAPIM clade (Fig. S75, Align. S106, Data S134). The removal of the joint MAPIM + *Asfarviridae* clade (Fig. S76, Align. S107, Data S135) was reciprocal since *Lavidaviridae* and *Herpesviridae* were added to the alignment S27. The resulting topology showed that the position of herpesviruses and virophages remained unchanged, which implies the absence of the homoplasies. Thus, the SFIII helicases, along with PolBs, D5-like primases, and D6/D11 helicases, were the fourth protein (gene) of the *Adenoviridae*/Polintons replication module, presumably acquired from within NCLDV.

### The transcription and RNA processing proteins

Reconstruction of the mRNA capping enzyme tree, consisting of the eukaryotic, NCLDV, and CLP homologs, showed the origin of the plasmid sequences from within NCLDV, which was consistent with the data on PolB, and D6/D11 helicases (Fig. S77, Align. S108, Data S136). The removal of the sequences of marseilleviruses, pithoviruses, and cedratviruses from the alignment did not affect the topological position of the CLP clade (Fig. S78, Align. S109, Data S137). This suggested that, like the PolB and D6/D11 genes, the ancestral mRNA capping enzyme gene of CLPs presumably also was acquired from within NCLDV.

We got a different result for the RNAP second largest (β) subunits of CLPs compared to phylogenies of PolB, D6/D11, and mRNA capping enzyme. The CLP clade was found localized sisterly to the RPA2 clade (Fig. S79, Align. S110, Data S138). Removal of the long RPA2 branch sequences from the alignment S110 did not change the topological position of the CLP clade (Fig. S80, Align. S111, Data S 139). To verify the result, we aligned the crenarchaeal, eukaryotic, and plasmid sequences using the MUSCLE v5 software and reconstructed the RNAP second largest (β) subunit tree. The resulting topology confirmed the sister relationship between the plasmid and RPA2 clades (Align. S112, Data S140). Considering the sister relationship between the NCLDV and RPB2 clades, as well as the sister relationship between the CLP and RPA2 clades, we rejected the possibility of the origin of the ancestral second largest (β) subunit gene of CLPs from within NCLDV.

The interim conclusion from our data on the genome replication and expression machinery suggests that several genes of the *Adenoviridae*/Polintons lineage (except for the RNAP second largest (β) subunit of CLPs) and herpesviruses, originated from within the NCLDV clade.

### The morphogenetic module

There are conflicting data in the literature on the belonging of adenoviral IVA2 to different branches of ASCE ATPases (52, 53). It has been shown that the adenoviral packaging ATPases and terminases have an additional strand to the “right” of the core P-loop domain, which suggests their evolutionary relationship with the FtsK-HerA and PilT/VirB11 superfamilies (52, 60). At the same time, it has been proposed that adenoviral IVA2 ATPases may have originated from the ABC superfamily ATPases (53). IVA2 and ABC ATPases share two polar residues at the end of the sensor-1 strand, one of which is highly conserved histidine (53). In addition, the prediction of the secondary structure of IVA2 suggests that they contain an insert with β-strands after helix-1 (53). However, ABC ATPases after β-strands contain one more strand, which distinguishes them from IVA2 (52, 53). We aligned the Helix-1 – Strand 4 (sGsGKo – Q) fragment of the IVA2, HerA, and FtsK sequences with the bacterial and eukaryotic ABC homologs and reconstructed a tree. We proceeded from the assumption that if the adenoviral Helix-1 – Strand 4 fragment potentially originated from the ABC transporters, then the adenoviral Helix-1 – Strand 4 clade must be localized outside of the FtsK-HerA clade. However, the resulting topology nested the *Adenoviridae* Helix-1 – Strand 4 fragment in the HerA-FtsK clade (Fig. S81, Align. S113, Data S141). Localization of the Helix-1 – Strand 4 clade of adenoviruses actualized the construction of a tree consisting of the *Adenoviridae*/Polintons lineage, as well as their cellular homologs (Fig. S82, Align. S114, Data S142). At this step, we also used the anchor amino acid substitutions belonging to the Helix-1 – Strand 4 (sGsGKo – Q) fragment highlighted in Figure 2 (52). The resulting topology showed that the *Adintoviridae*/Polintons clade was a sister to the FtsK clade. The addition to the analysis of the A32-like, *Pleurochrysis sp. endemic virus 2*, *Pleurochrysis sp. endemic virus 1a*, *Bovine rumen MELD virus*, and *Sheep rumen MELD virus* homologs showed that the *Adenoviridae*/Polintons clade nested within NCLDV (Fig. 15, Align. S115, Data S143). Such a topological localization of the *Adenoviridae*/Polintons clade was consistent with the PolB and SFIII helicase data (see above). To verify the potential influence of homoplasies on the topological position of the *Adenoviridae*/Polintons clade, we a) removed the sister mimiviruses from the alignment S115, b) removed the entire outgroup of the joint NCLDV + *Adenoviridae*/Polintons clade from the alignment S115 and reconstructed trees (Figs. S83-S84, Aligns S116-S117, Data S144-S145). The results confirmed the origin of the FtsK-HerA ATPases of the *Adenoviridae*/Polintons clade from within the joint *Mimiviridae* + MAPIM clade. However, the moderate and low statistical supports of the *Mimiviridae* + *Adenoviridae*/Polintons nodes (see Fig. 15 and Fig. S84) did not allow confident identification of a sister relationship between the *Adenoviridae*/Polintons and *Mimiviridae* clades.

**Fig. 15.**
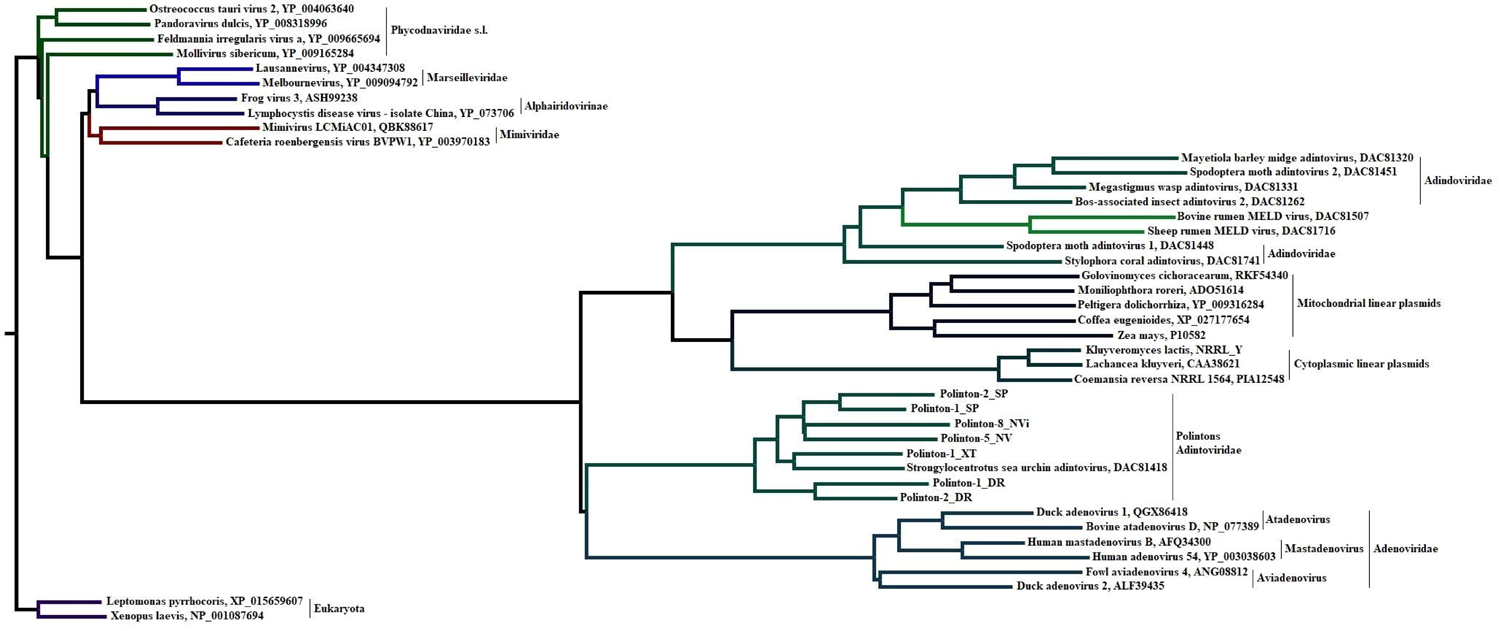
Maximum likelihood phylogenetic tree of the FtsK-HerA superfamily ATPases. The tree consists of the archaeal HerA, bacterial FtsK, ViB3, VirB4, A32-like (NCLDV), IVA2 (*Adenoviridae*), Polintons, *Adintoviridae*, *Lavidaviridae*, *Pleurochrysis sp. endemic virus 2*, *Pleurochrysis sp. endemic virus 1a*, *Bovine rumen MELD virus*, and *Sheep rumen MELD virus* homologs. The tree was constructed using the IQ TREE program (version 1.6.12) under the EHO+I+R4 model and rooted according to the topology of Fig. 6, as well as in published data (52). The corresponding alignment was built using the MUSCLE algorithm. Statistical support of the partitions, measured by SH-aLRT branch tests (1,000 replicates), approximate Bayes, and ultrafast bootstrap (1,000 replicates), is shown at the nodes. Scale bars represent the number of amino acid substitutions per site.

To identify phylogenetic relationships of the DJR MCP proteins of the NCLDV clades and *Adenoviridae*/Polintons lineage members, we built two alternative alignments, consisting of the NCLDV, *Adenoviridae*, *Lavidaviridae*, *Adintoviridae*, *Yaravirus brasiliensis*, *Pleurochrysis sp. endemic virus 1a*, *Pleurochrysis sp. endemic virus 1b*, *Pleurochrysis sp. endemic virus 2*, *Bovine rumen MELD virus*, *Sheep rumen MELD virus*, and Polintons sequences. To create the first alignment, the sequences of the *Adenoviridae*/Polintons lineage including the listed unclassified viruses were aligned first, and then that alignment was aligned with the prealigned NCLDV sequences (Fig. S85, Align. S118, Data S146). For the second alignment, the sequences of the *Adenoviridae*/Polintons lineage including the unclassified viruses were aligned within families/clades frameworks, and then we aligned these separately prealigned blocks with the prealigned NCLDV sequences (Fig. S86, Align. S119, Data S147). In both cases, the *Adenoviridae*/Polintons lineage formed a clade, which spoke in favor of the monophyletic origin of their DJR MCPs. However, in the case of the alignment S118, the *Adenoviridae*/Polintons clade was localized sisterly to mimiviruses. Meanwhile, topology derived from the alignment S119 localized the *Adenoviridae*/Polintons clade sisterly to iridoviruses. We removed the corresponding sister clades from both alignments (Aligns. S118-S119) and reconstructed trees (without *Mimiviridae* – Fig. S87, Align, S120, Data S148; without *Iridoviridae* – Fig. S88, Align. S121, Data S149). In the case illustrated in Figure S85, the removal of mimiviruses resulted in a change in the expected position of the *Adenoviridae*/Polintons clade (Fig. S87). In contrast, in the case illustrated in Figure S86, the removal of iridoviruses did not lead to a change in the expected position of the *Adenoviridae*/Polintons clade (Fig. S88). Thus, according to the formal criterion, we preferred the alignment S119 and its derivative topology, since the alignment S118 exhibited some level of homoplasies. Meanwhile, it should be noted that the position of the AP clade in Figure S86 corresponded only to the topology of the dendrogram shown in Figure 5.

Next, based on the pairwise Z score comparisons, we calculated the dendrogram of the full-length DJR MCP proteins of NCLDV, *Lavidaviridae*, *Adenoviridae*, and Polintons using the I-TASSER predictions and crystallographic structures deposited in the PDB (Fig. 16, Data S150). The *Adenoviridae*/Polintons clade, consisting of the PDB crystallographic structures and I-TASSER predictions, formed a sister clade to the MAPIM clade which spoke in favor of the topology shown in Figure S86 (Align. S119). We also note that dendrograms calculated using only the I-TASSER predictions (Data S151) or using only the PDB crystallographic structures (Data S152) gave the same direction of evolution as the combined dendrogram shown in Figure 16.

**Fig. 16.**
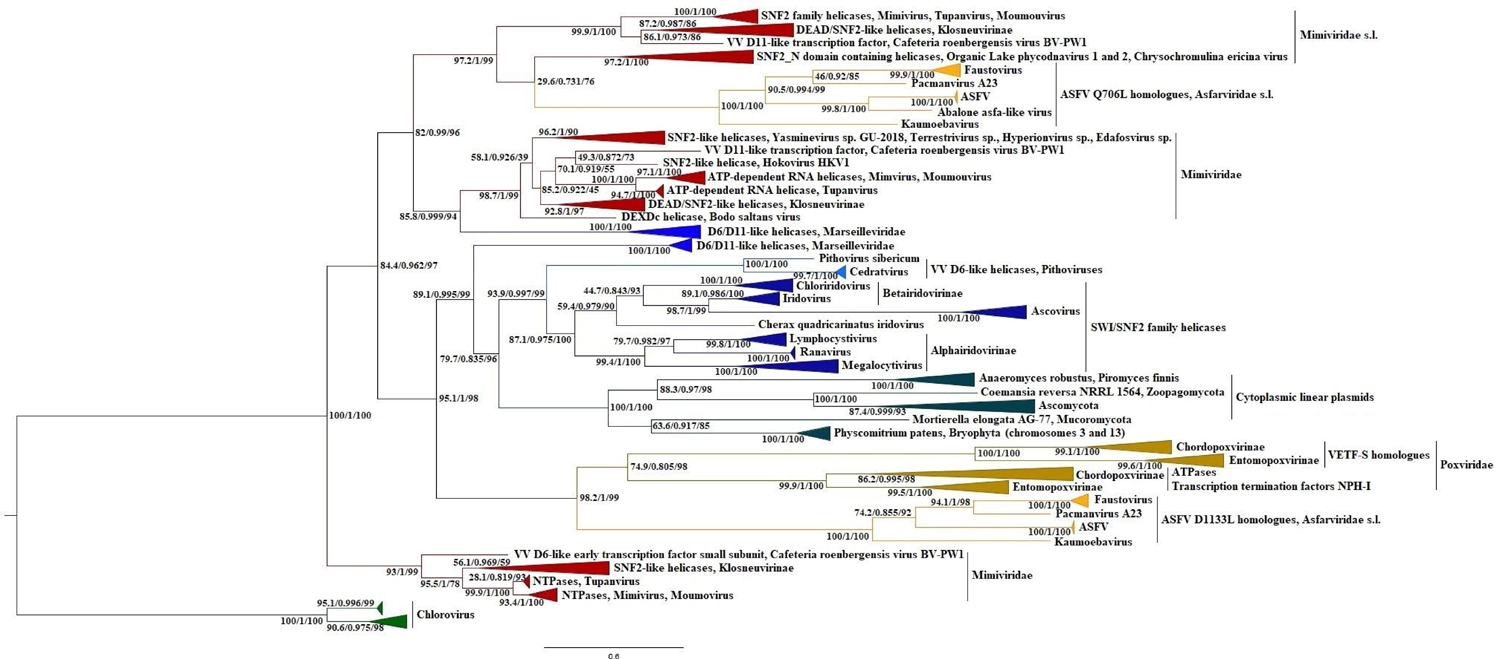
The NCLDV, *Lavidaviridae*, and *Adenoviridae* DJR MCPs dendrogram. The dendrogram is based on the pairwise Z score comparisons calculated using DALI. Protein structure prediction was carried out using I-TASSER (54–56). The dendrogram was rooted between *Phycodnaviridae* and the rest of the NCLDVs according to the topologies of all individual phylogenies containing the cellular and NCLDV clades presented in this work. GenBank accessions are given for the sequences used for the I-TASSER predictions, as well as for the crystallographic structures deposited in the PDB. The I-TASSER prediction was carried out without removing any amino acid residue within the sequences.

## Discussion

The taxonomy of *Bamfordvirae* should be based on the identified branching order of monophyletic clades that form this kingdom. In the context of this idea, NCLDV is a good entry point into unraveling the branching order of *Bamfordvirae* since genomes of NCLDV contain more than a dozen of nearly-omnipresent core genes that have homologs in eukaryotes and bacteria which can be used as outgroups (2, 7–8).

Most of the core genes presumably had been transferred from a lineage leading to modern eukaryotes into the genome of the LCA of NСLDV. Alternatively, they potentially had been acquired from various now-extinct protoeukaryotic lineages. In addition, several core genes are of bacterial origin and their evolutionary path in terms of now-existing or now-extinct bacterial lineages could be similar to the evolutionary path of core genes of protoeukaryotic origin. Gene acquisition events could have taken place many times and from different sources. Therefore, in the present day, they can be a source of a heterogenic plesiomorphic set of signals. Potential heterogeneity of NCLDV plesiomorphic signals can increase the likelihood of homoplasies between the clades of modern NCLDVs and the eukaryotic or bacterial outgroups, especially given the more rapid accumulation of substitutions in viral lines (long branches). There are two possibilities here. We could get conflicting evolutionary trajectories of NCLDV for the phylogenies and dendrograms of various individual proteins, as well as a unified one. However, we confidently obtained the realization of the second probability, that is, all phylogenies and dendrograms of individual core proteins of NCLDV gave the same direction of evolution. This means that in the conflict between synapomorphies and homoplasies, the contribution of synapomorphic signals of NCLDV for the reconstruction of the evolutionary trajectory of this group of giant viruses proved to be critically decisive.

The consistency and repeatability of our results were due, among other things, to the taxonomically widest use of outgroups (37, 61). To combat potential homoplasies affecting the topological relationships among NCLDV clades we applied several approaches identifying the long branch attraction effect as well as approaches that eliminate this effect described in the literature (45). Firstly, 15 individual protein phylogenies and 10 individual protein dendrograms where outgroups were used showed the same direction of evolution of NCLDV for major clades (first bifurcation between the *Phycodnaviridae* s.l. and the MAPIMAPM clades). Such unanimity spoke of the correctness of the found direction of evolution of NCLDV. Except for PolB and Metallopeptidase WLM phylogenies, all other individual phylogenies and dendrograms also supported the second bifurcation between the MAPIMAP and *Mimiviridae* s.l. clades. This finding allowed us to suggest that the topological position of the AP clade polymerases among NCLDVs could be due to homoplasies. However, careful analysis, including the calculation and construction of various dendrograms and phylogenies, as well as morphological analysis of the beta-hairpin predictions, spoke in favor of the relationship of the AP clade polymerases with the δ catalytic subunits and NCLDV polymerases but did not support affinity with polymerases of baculoviruses, nudiviruses, nimaviruses, malacoherpesviruses, and alloherpesviruses (22). Nevertheless, the position of the AP clade polymerases awaits further study. Secondly, we built supermatrixes consisting of core NCLDV proteins of protoeukaryotic and bacterial origin. A comparative analysis of the seven concatenated topologies did not reveal the influence of outgroups on the branching order within NCLDV. The same direction of evolution of NCLDV and relationships among clades within NCLDV were obtained via the construction of the supertree and superdendrogram. It is precisely the concatenated trees, supertree, and superdendrogram that have expressed a preference for sister relationships between the AP and MAPIM clades. Thirdly, the same phylogenetic relationships between the AP and MAPIM clades, as well as between the MAPIMAP and *Mimiviridae* s.l. clades were obtained using extremely closely related viral homologs of the D6/D11 helicases as an outgroup. Fourth, topologies of outgroups were examined separately before being combined with NCLDV sequences. Outgroups were checked for the identification of horizontal transfers and homoplasies, that is, for the preservation of monophyly of clades constituting outgroup, as well as for compliance with the phylogenies given in the literature. Fifth, to test the stability of the topological relationships among the NCLDV clades of individual protein trees, we changed the composition of outgroups, as well as constructed the same trees without outgroups (35, 45). Sixth, to check the topological position of any clade, we used reciprocal deletion of clades, even if only one clade was long. In this case, we proceeded from the assumption that if the sequences of the only single clade formed a long branch, i.e., had an increased number of amino acid substitutions per site compared to the sequences of a neighboring branch, this could become a source of attraction and lead to the artificial topology.

Despite these efforts, we found topological differences not related to the first bifurcation event between individual phylogenies and individual dendrograms of the same proteins. We suggest to the reader two possible explanations for this phenomenon. The most likely explanation is the use of bioinformatics analysis of the evolutionary history of proteins to predict the tertiary structures of proteins (46–48). The second potential explanation is more speculative. It is possible that within certain taxonomic units, in our case within the *Nucleocytoviricota*, such elements of protein secondary structures as β-sheets and α-helices have a spatial framework in which they can change their spatial position. In other words, in the course of evolution, an element of the protein architecture belonging to some crown clade can change its spatial position, returning to a plesiomorphic spatial position, which can hypothetically affect the topology of the dendrogram. Regardless of the reasons, we briefly repeat the thought expressed above. At long evolutionary distances, the calculation of dendrograms must be accompanied by the construction of phylogenies and the study of protein morphologies.

Our study revealed that according to the vast majority of the individual phylogenies and dendrograms, as well as supertree, superdendrogram, and concatenated trees phycodnaviruses s.l. resolved as a sister to the MAPIMAPM clade. The exceptions were the phylogenies of SFIII helicases and Uracil DNA glycosylases. Proteins of phycodnaviruses s.l. were separated from NCLDVs by the bacterial or prophage homologs in these cases. However, the gene was replaced secondarily in the case of the SFIII helicases in phycodnaviruses s.l., or vice versa, in the LCA of MAPIMAPM. The gene was found only in pandoraviruses from among phycodnaviruses s.l. in the case of Uracil DNA glycosylase. The product of this gene was localized among eukaryotic and bacterial homologs, which indicates a separate fate of the *Pandoraviridae* homologs from the rest of the NCLDVs. The scarcity of alternative scenarios suggests that the first bifurcation of the LCA of NCLDV must have led to the emergence of two branches. The first branch led to *Phycodnaviridae* s.l. including pandoraviruses, molliviruses, and medusaviruses. The second branch led to the LCA of the MAPIMAPM clade, which, in turn, branched into the *Mimiviridae* s.l. and the MAPIMAP clades (Figs. 8-10). This branching order contradicts the accepted ICTV taxonomy based on the data presented in the literature (2, 7, 9-10). The taxonomy of ICTV is fully consistent with the branching order of NCLDV presented in 2009 and 2019 (7, 10). Koonin and Yutin gave arguments justifying the branching order of the NCLDV, according to which NCLDV consists of three clades, i.e., *Mimiviridae* + *Phycodnaviridae* (branch 1), MAPIM (branch 2), and AP (branch 3) (10). Their concatenated tree was constructed without using cellular xenologues and technically was unrooted. The choice of branching order was based on two arguments - the greater similarity of the core protein sequences of branches 1 and 2 (combined into the class *Megaviricetes*) in comparison with branch 3 (the class *Pokkesviricetes*) and on the topologies of the phylogenetic trees of individual proteins, in particular, helicase-primase, as well as DNA and RNA polymerases, although individual protein trees gave conflicting topologies (8). Phylogenies presented by other authors’ also included only viral sequences and were technically unrooted (9, 11, 43). Our phylogenetic trees obtained using cellular outgroups refutes the most important argument about the greater similarity of the proteins of branches 1 and 2 in comparison with homologs of branch 3. (10). Moreover, despite using bioinformatics data on the evolutionary history of proteins (ColabFold predictions), our dendrograms and superdendrogram also confirmed the similarity of the tertiary structures of the AP and MAPIM clades (branches 3 and 2 respectively according to Koonin and Yutin). Thereby, according to our results, the class *Pokkesviricetes* should be abolished, and the families *Asfarviridae* s.l. and *Poxviridae* should be included in the class *Megaviricetes* to reconcile phylogeny and taxonomy.

The monophyletic origin of the *Adenoviridae*/*Lavidaviridae*/*Adintoviridae*/CLP/MLP/Polintons lineage was supported by the phylogenies of the replication module proteins, as well as the mRNA capping enzyme and morphogenetic module proteins. Moreover, all these phylogenies and dendrograms, except for the RNAP second largest (β) subunit phylogeny, localized the *Adenoviridae*/Polintons lineage as a clade within NCLDV. The formation of the FtsK-HerA superfamily ATPase, DJR MCP, D5-like primase, D6/D11 helicase, SFIII helicase, PolB and mRNA capping enzyme clades by the *Adenoviridae*/Polintons lineage and localization of these clades within NCLDV casts doubt on the monophyly of *Preplasmiviricota* and suggests that the viruses of the *Adenoviridae*/Polintons lineage should be placed within the *Nucleocytoviricota* phylum.

The CLP and MLP clades localization within NCLDV suggests that unlike the strategies of vertical inheritance of genetic information (viruses) and integration into the host genome (Polintons) the ancestor/s of CLPs and MLPs could be a virus related to NCLDV, which chose the third, intermediate strategy - life in the cytoplasm and mitochondrial matrix.

As we have shown, the PolB, SFIII helicase, and ASFV F1055L homologs of herpesviruses presumably originated from within NCLDV. A possible explanation for these findings may be the assumption of simultaneous infection of eukaryotic cells with NCLDV and ancient herpesviruses, which could result in the horizontal transfer of the replication module genes from NCLDV to herpesviruses. However, some differences in the topological localization of herpesvirus proteins within NCLDV likely indicate that these hypothetical horizontal transfers occurred more than once, and gene donors belonged to different clades of NCLDV.

Four competing scenarios have been proposed to explain the origin of viruses. The Coevolution hypothesis (Bubble Theory) postulates the origin of viruses from precellular genetic elements (62). According to the second hypothesis (‘regression’ or ‘reduction’ hypothesis), viruses evolved from cellular life forms via the reduction of genomes. The third hypothesis involves the escape of genes from cellular hosts and the acquisition of partial replicative autonomy. The fourth “chimeric” hypothesis has been recently proposed and suggests that the replicative modules of different viral lineages may be of primordial, precellular origin, while capsid proteins have been captured from cellular organisms in different evolutionary periods multiple times and independently (6, 63). There also may be combinations of some or all of those four scenarios. Moreover, the first and “chimeric” hypotheses rather complement and enrich each other than contradict. Our results do not refute the “chimeric” hypothesis about the primordial origin of the replication module, in particular, for monophyletic *Riboviria* (64). It is also possible that the substantial portions of replicative modules of *Bamfordvirae* and *Herpesviridae* were of primordial origin. However, it is possible that some of them were replaced with the corresponding host genes, as in the case of *Bamfordvirae*, where we observe the presence of eukaryotic xenologues. As shown above, apparently three genes of the replicative module in *Herpesviridae* originated from within NCLDV. Likewise, one of the key genes of the morphogenetic module of *Bamfordvirae* the FtsK-HerA superfamily ATPases was recruited from bacterial homologs.

Finally, the phylogenies and dendrograms of the *Adenoviridae*/Polintons lineage presented and discussed here disproved the possibility of the origin of NCLDV from the *Adintoviridae*/Polinton-like viruses (4, 5) and vice versa, our data supported the hypothesis of the origin of NCLDV from prokaryotic viruses with small genomes proposed by Yutin and colleagues (3).

## Materials and Methods

### General information

In this study, we used 18 core proteins of NCLDV, as defined by Yutin and colleagues and Aylward and colleagues (7, 43). Correspondences of the names of the used proteins, NCVOGs and GVOGs are given in Table S5. These proteins, together with their homologs from the *Preplasmiviricota* phylum, were collected from genomes belonging (or tentatively assigned) to realm *Varidnaviria* and related mobile elements, as well as from genomes of *Herpesvirales*, *Nudiviridae*, *Baculoviridae*, *Nimaviridae*, and *Hytrosaviridae* (Table S6).

### Sources of amino acid sequences and gene prediction

Amino acid sequences were downloaded from the NCBI non-redundant protein sequences (nr) database (65). The genomes of Polintons were downloaded from the RepBase database (66, 67). The selection of Polintons genomes was made from the 20181026 release. The prediction of the Polintons genes has been carried out using the Heuristic Approach for Gene Prediction web server available at http://exon.gatech.edu/GeneMark/heuristic_gmhmmp.cgi (68, 69). For further work, we selected Polintons, whose genomes encode the PolB, FtsK-HerA superfamily ATPase, and DJR MCP genes or at least PolB and one of the latter two genes (Table S7).

### Sequence selection

In most families and genera of NCLDV, the majority of the core NCVOG (GVOG) representatives are single-copy (see Table S6 and 70). We used single-copy homologs of NCLDVs as queries and performed the blastp and PSI-BLAST searches against the nr and env_nr databases (71, 72). For each search, we entered taxon ID and downloaded only subjects with the highest max score for each species (E value < 0.005). Then, the same procedure was carried out with the received objects, thus maximally covering the entire possible range of subjects.

The initial pool of cellular homologs was selected by blastp and PSI-BLAST searches against the nr database where queries were the NCLDV sequences. We selected the cellular subjects with the highest max scores and used them as queries as described for viral sequences.

After the accumulation of a redundant pool of viral and cellular homologs, we constructed the LG (+ accessories) preliminary trees to remove horizontally transferred and redundant sequences from the alignments. See Text S1 for details.

We used as queries not only the sequences identified as a result of seeding of the NCLDV sequences but also sequences presented in the literature in the cases of DNA polymerases and FtsK-HerA superfamily ATPases.

The list of GenBank accessions used in this study is presented in Table S8.

### Multiple sequence alignments and multiple protein structure alignments

Multiple sequence alignments were generated using the MEGA7 software package (73) by the MUSCLE (74) algorithm. To verify the topologies of some key trees obtained using MUSCLE, we used the MAFFT online 7.47 version available at https://www.ebi.ac.uk/Tools/msa/mafft/ (75–76), as well as the MUSCLE v5 software (44). For the MUSCLE v5 software, we used the mpc algorithm and the permutation none, perturbation 0 options. The MUSCLE v5 software was also used to generate alignments S29 and S30 (cellular and viral SFII helicases). See Text S1 for details.

Alignments for concatenation were built based on the alignments of the corresponding individual proteins, with the addition/deletion of some sequences. Otherwise, they were treated the same as the rest of the alignments (see below). Head-to-tail linking of the aligned (MUSCLE) individual protein alignments was used to build the supermatrixes. In concatenated alignments, due to the absence of some proteins in some taxa, gaps were not removed.

Multiple protein structural alignments were performed on the mTM-align server available at https://yanglab.nankai.edu.cn/mTM-align/ (41, 42).

### Phylogenetic analysis

Best-fit general amino acid replacement matrix models were selected from the common model list of the IQ TREE program (version 1.6.12) (77) using the -m TESTONLY and -m TESTNEWONLY commands. The Bayesian selection criterion (BIC) was used by default. Single-protein and concatenated protein phylogenetic trees were constructed within the maximum likelihood (ML) framework using the IQ TREE program (version 1.6.12) (77). Sets of trees based on individual protein and concatenated protein alignments were constructed under the LG, LG4X, LG4M, LG+C20, EX2, EX3, EHO, EX_EHO, UL2, and UL3 (+ accessories) models. Exceptions were made for the PolB phylogenies constructed using a large number of metagenomic sequences (Fig. 2, Fig. 12, Data S8, Data S10), for the SFII helicases (Data S28-S29), and for checking the topological position of RNAP second largest (β) subunit of CLPs (Data S140). In these cases, the trees were constructed under the best-fit models (-m TESTONLY). For statistical support of the partitions on the trees obtained by IQ TREE, we used four tests, i.e., the SH-aLRT branch, approximate Bayes (aBayes), ultrafast bootstrap (Ultrafast), and standard bootstrap (78–82). The default parameters (1000 for SH-aLRT and Ultrafast) were applied, and the number of replicates for the standard bootstrap test was set to 100. Sets of trees obtained from all individual and concatenated protein alignments were subjected to topology testing using the TreeFinder program under the LG+G4 model (83). We applied the following tests: ELW, BP, SH, WSH, AU, and likelihood (78, 84–86). The results of topological tests of individual and concatenated proteins are shown in Tables S9-S28. The best AU or AU/likelihood trees were used for visualization and supertree construction.

### Supermatrix analysis

We obtained seven supermatrixes by concatenating the alignments, the protein compositions of which are shown in Table S4. Due to the limitations of the taxonomic composition of the alignments used for concatenation, each alignment was subjected to a verification procedure starting from the stage of construction of a preliminary LG (+ accessories) tree as described above and in Text S1. Any sequence violating the monophyly of another clade was removed from the further analysis. Information about concatenated trees construction, as well as topological tests, is given above.

### Supertree construction

To construct supertrees, we constructed the set of input bootstrapped trees for 13 protein families (23 trees in total) the substitution matrices of which corresponded to the best AU values (Table S26). To avoid obvious errors caused by horizontal gene transfers and homoplasies into supertrees, for the construction of input bootstrapped trees we used the same condition as in the case of construction of concatenated supermatrix, i.e., preservation of monophyly of well-established clades. We limited the taxonomic composition of the sequences used to 61 species (52 viral and 9 eukaryotic). The supertree was constructed using the subtree prune-and-regraft distance (SPR supertrees) method. We used the SPR Supertree 1.2.1 downloaded version (58). We constructed a supertree with bootstrap thresholds equal to 80 and the -allow_multi and -i 1000 options. The rest of the options were used by default.

### Identification of the PolB metagenomic sequences

The PolB metagenomic sequences (43) were identified using HHPred screening (87–88) available at https://toolkit.tuebingen.mpg.de/ in the PDB_mmCIF70_14_Apr, ECOD_F70_20220113, UniProt-SwissProt-viral70_3_Nov_2021, NCBI_Conserved_Domains(CD)_v3.19 databases. To identify these sequences, we also performed a PSI-BLAST screening in the nr database.

### Determination of the root position

The root position of individual phylogenies, dendrograms, concatenated trees, supertree, and superdendrogram was determined by the presence of cellular and viral (D6/D11 helicase) homologs used as outgroup (see Text S1 for details). The root position of phylogenies, where the cellular outgroups were not used, was determined according to the direction of evolution of NCLDV obtained for the same proteins whose trees were constructed using the cellular outgroups. For “synapomorphic” proteins, the root position was determined according to the direction of evolution of NCLDV identified for all individual phylogenies.

### Prediction of three-dimensional structures of proteins using ColabFold

Except for the DJR MCP sequences, the sequences of interest were aligned via the MUSCLE v5 software using the mpc algorithm and the permutation none, perturbation 0 options. To exclude spatial fluctuations that are not typical for most proteins of a given alignment and that could potentially affect the results of the analysis we removed from the alignment fragments between constant blocks, formed by single or/and unaligned sequences. Thus, we obtained presumably functionally active standardized fragments of the minimum size, from 80 to 650 amino acids in length.

To predict the three-dimensional structures of protein fragments, we used the online version of the ColabFold software, which speed-ups the three-dimensional prediction of protein structures by replacing the AlphaFold2’s input feature generation stage with a fast MMseqs2 available at https://colab.research.google.com/github/sokrypton/ColabFold/blob/main/AlphaFold2.ipynb (46–48, 89). The options were used by default.

For dendrograms calculation and the Exo I – MotifC fragments visualization, we selected from the redundant set of predictions a subset of predictions consisting of predictions of the unrelaxed rank 1 with the pLDDT > 70 for an entire prediction. The length and quality of predicted fragments were controlled using the output predicted aligned error and predicted LDDT per position plots. See Text S1 for details.

Used in this work the unrelaxed rank 1 predictions (.PDB files), corresponding GenBank accessions, corresponding RepBase codes of Polintons, as well as alignments (.fas) of predicted fragments, are given in (Data S153-S419 (predictions), Data S420-S430 (alignments)).

### Prediction of the three-dimensional structures of the DJR MCPs proteins using I-TASSER

We used a threading method implemented in the I-TASSER (Iterative Threading ASSEmbly Refinement) program for full-size DJR MCP structures prediction available at https://zhanggroup.org/I-TASSER/about.html (54–55). Predicted structures, as well as corresponding GenBank accessions, are given in Data S431-438.

### Dendrograms and superdendrogram calculation

Structural comparisons of the ColabFold and I-TASSER predictions with the crystallographic structures deposited in the PDB were performed using the PDB25 algorithm via the DALI server (49–51). Exhaustive PDB25 search results of the ColabFold and I-TASSER predictions are shown in Table S3. The ColabFold and I-TASSER predictions intended for dendrograms calculation had Z-scores > 2 relative to corresponding subjects (49–50). Dendrograms were calculated using the DALI server. Z scores of calculated dendrograms are shown in Table S1.

Based on 12 ColabFold-derived dendrograms (Data S439-S450) of the 9 core NCLDV proteins we constructed the SPR superdendrogram using the SPR Supertree 1.2.1 downloaded version (58). For calculation of a superdendrogram we used the -reroot and -i 10000 options. The rest of the options were used by default.

### Visualization

The phylogenetic trees, dendrograms, concatenated trees, supertree, and superdendrogram were rooted and visualized via FigTree v1.4.4 available at http://tree.bio.ed.ac.uk/software/figtree/. The Family B polymerase predictions (Fig. 4) were visualized via The PyMOL Molecular Graphics System, Version 4.6, Schrödinger, LLC.

## Acknowledgments

We are grateful to Dr. Arcady Mushegian and Dr. Darius Kazlauskas for many helpful discussions and advice in the course of this project.

## Data availability

The supporting files for this article (Figs. S, Align. S, Data S, and Tables S) can be found online at: https://zenodo.org/record/6790805#.YsBWinZByUk DOI 10.5281/zenodo.6790428

---

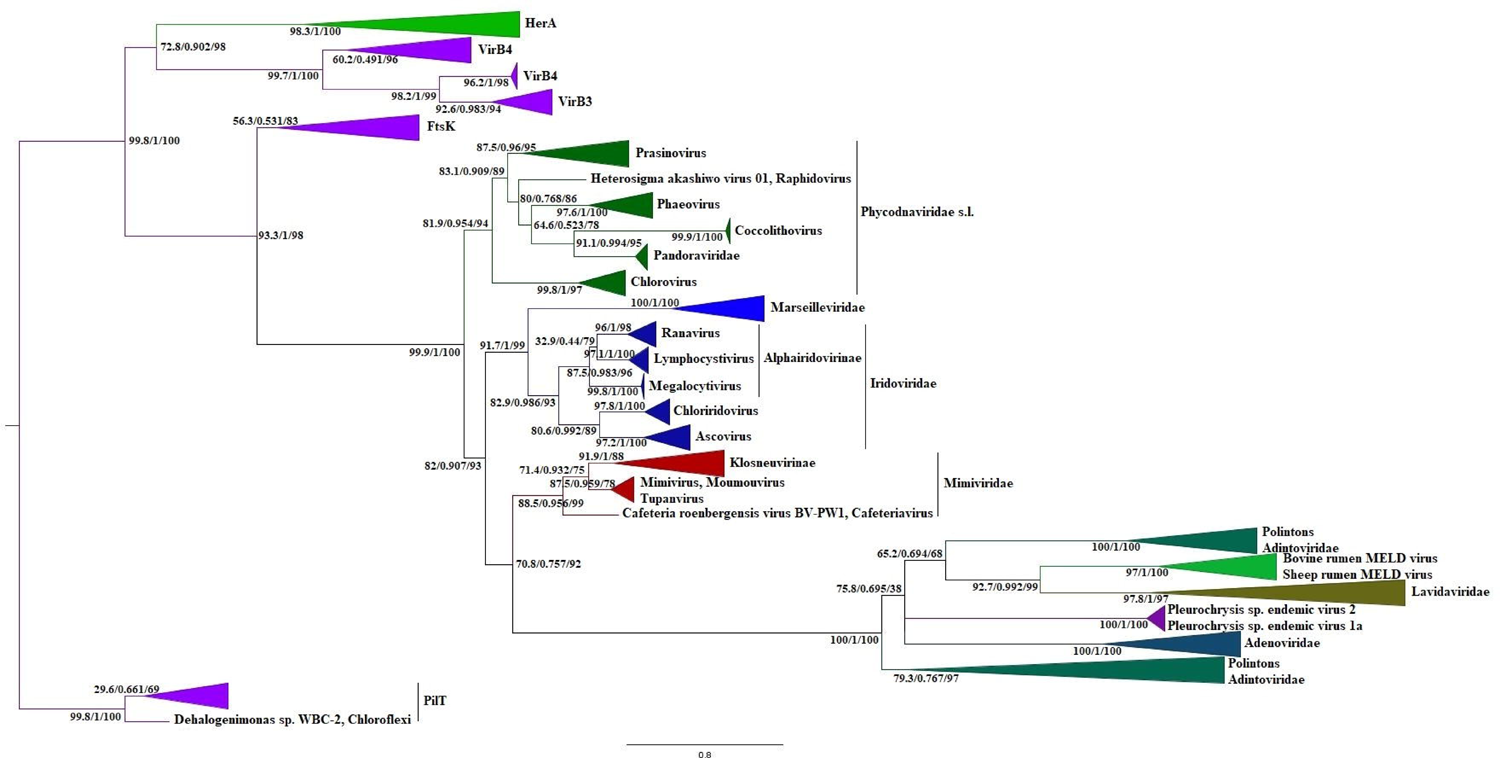

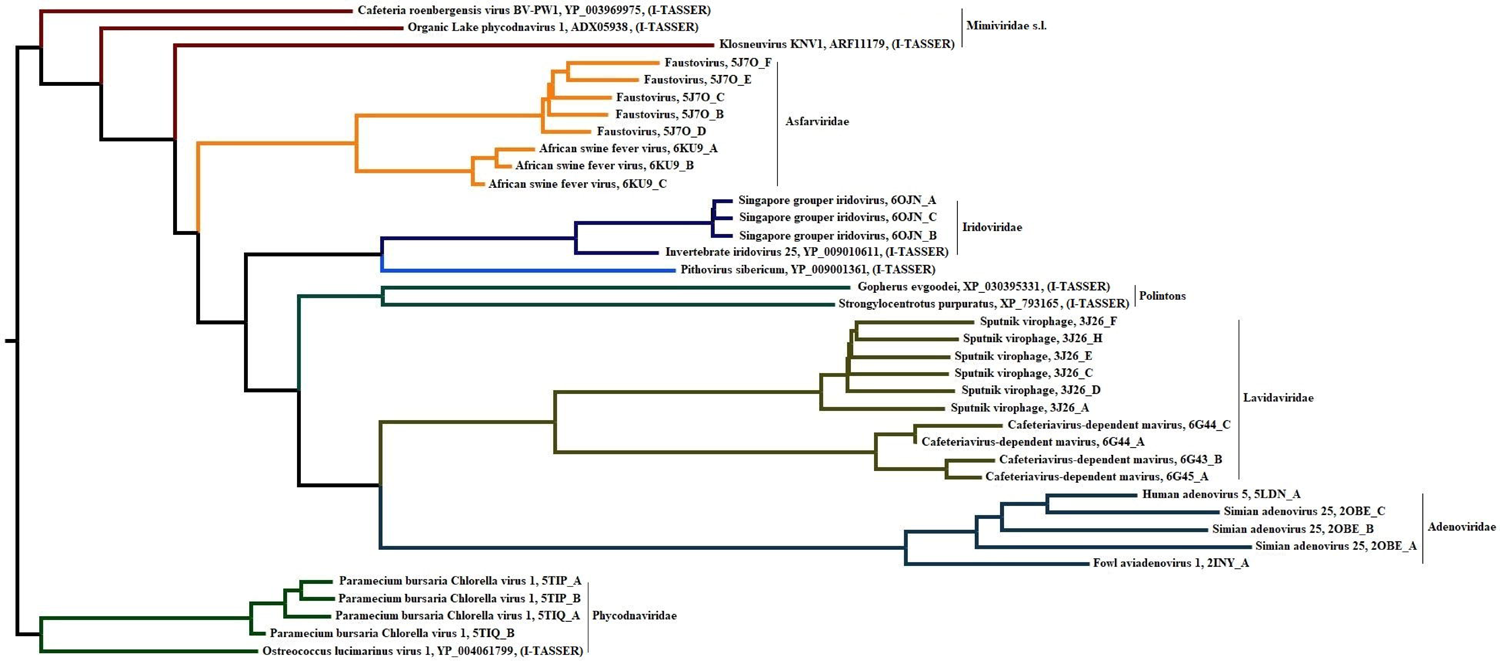

